# Postnatal FGFR-signaling establishes gradients of secretory cell identities along the proximal-distal axis of the lung airways

**DOI:** 10.1101/2023.12.11.571142

**Authors:** Alexandros Sountoulidis, Alexandra B. Firsova, Andreas Liontos, Jonas Theelke, Janine Koepke, Pamela Millar-Büchner, Louise Mannerås-Holm, Åsa Björklund, Athanasios Fysikopoulos, Konstantin Gaengel, Fredrik Bäckhed, Christer Betsholtz, Werner Seeger, Saverio Bellusci, Christos Samakovlis

## Abstract

Secretory cells are major structural and functional constituents of the lung airways. Their spatial organization and specification mechanisms are partially understood. Here, we labelled major secretory airway cell types and analysed them at single-cell resolution. We found opposing, partially overlapping gene-expression gradients along the proximal-distal airway axis superimposed on a general gene program encoding detoxification. One graded program is elevated proximally and relates to innate immunity, whereas the other is enriched distally, encoding lipid metabolism and antigen presentation. Intermediately positioned cells express low levels of both graded programs and show increased clonogenic potency in vitro, relating cell-plasticity to location in each branch. Single-cell RNA-sequencing following lineage-tracing revealed the sequential and postnatal establishment of the gradients in common epithelial progenitors. Fgfr2b is distally enriched and its postnatal inactivation reduces distal gene expression and expands proximal genes into distally located cells. This suggests a central role of FGFR-signaling in tissue-scale airway patterning.

## Introduction

The airway epithelium functions as a pathogen barrier and as a seamless conduit of air to the alveoli, where gas exchange takes place^1^. The airway network is anatomically divided into the extra-lobar (trachea and bronchi) and intra-lobar compartments^2^ with distinct cell compositions. Basal cells, for example, are localized in the extra-lobar compartment of the mouse airways^3^ and gradually decrease along the proximal-distal (PD) axis of the human intra-lobar airways^4^. Advances in single-cell RNA Sequencing (scRNA-Seq) resulted in the detailed characterization of the epithelial cell heterogeneity in mouse and human trachea, capturing the gene expression profiles of many cell types with single cell resolution^5–7^. It was only recently shown that specific cell types expressing secretoglobins, mucins and surfactant proteins are differentially distributed in distinct regions of human small airways^7^, even though pioneering experiments have reported differential expression for a few of these markers two decades ago^8^. The various epithelial cell types, distributed along the airways, are coordinated to accomplish distinct functions, creating the mucociliary escalator^9^. Secretory cells contribute to respiratory homeostasis by detoxification of inhaled xenobiotics^10^ and by secretion of mucins, antimicrobial proteins and cytokines upon exposure to pathogens^11–13^ ^8, 14^. Mucus and inhaled particles are propelled out of the airways by numerous multiciliated cells^15–17^. In homeostasis, secretory club cells self-renew and produce ciliated cells^18, 19^. Upon injury they can reconstitute the airways and even the alveolar epithelium^9, 20^. Several subsets of airway secretory cells contribute to tissue repair upon airway or alveolar injury. These include the variant (v) club cells^21, 22^, the *Upk3a^pos^* (u) club cells^23^ located near neuroendocrine cells (NE), the bronchioalveolar stem cells (BASCs) in terminal bronchioles (TBs)^22, 24, 25^, the β4^pos^ CD200^pos^ Scgb1a1^pos^ cells in distal airways^20^ and cells in a activated transitional states (ADI^26^, DAPT^27^ and PATS^28^). The existence of such a large variety of airway secretory cell states indicates that they are highly heterogeneous and capable of adopting distinct stem cell characteristics. Several of the described lung secretory cell types emerge postnatally, but their spatial coordinates in the tissue, lineage relationships and differentiation mechanisms are poorly understood.

We focused on the branched airway secretory epithelium in homeostasis and during development and captured the gene expression profiles and cell topology. Our work reveals a general airway secretory gene expression program in addition to at least two complementary and partially overlapping ones. These programs are genetically determined, established postnatally in an initial embryonic progenitor and relate to cell-topology along the branch PD-axis to define distinct functional characteristics, which are maintained in vitro in different types of secretory cells. We show that FGF-signalling in the distal branches is a major determinant of the spatial patterning of the airways, controlling genes involved in surfactant biosynthesis, mitochondrial function, ribosomal biogenesis and immune responses. Our study provides a detailed characterization of the lung secretory epithelium in time and space, suggests topology-related cell functions and provides mechanistic information on the dynamic nature of their regulation.

## Results

### Analysis of epithelial cell heterogeneity in the adult lung

To analyse the main secretory cell types in the mouse lung, we first isolated them from the lungs of a double transgenic reporter strain expressing a green fluorescence protein (GFP) in the alveolar secretory epithelium (Sftpc-GFP)^29^ and a tamoxifen-inducible red fluorescence protein (tdTomato) ^30^ in the airway secretory cells^19^ (Scgb1a1creER-Ai14). In addition, we used Pdpn antibodies in conjunction with the tdTomato fluorescence to label and isolate a third fraction because we had noticed that the cell surface protein Pdpn is expressed in basal cells and in secretory cells of the proximal airways^31^ (Fig. 1A).

**Figure 1.**
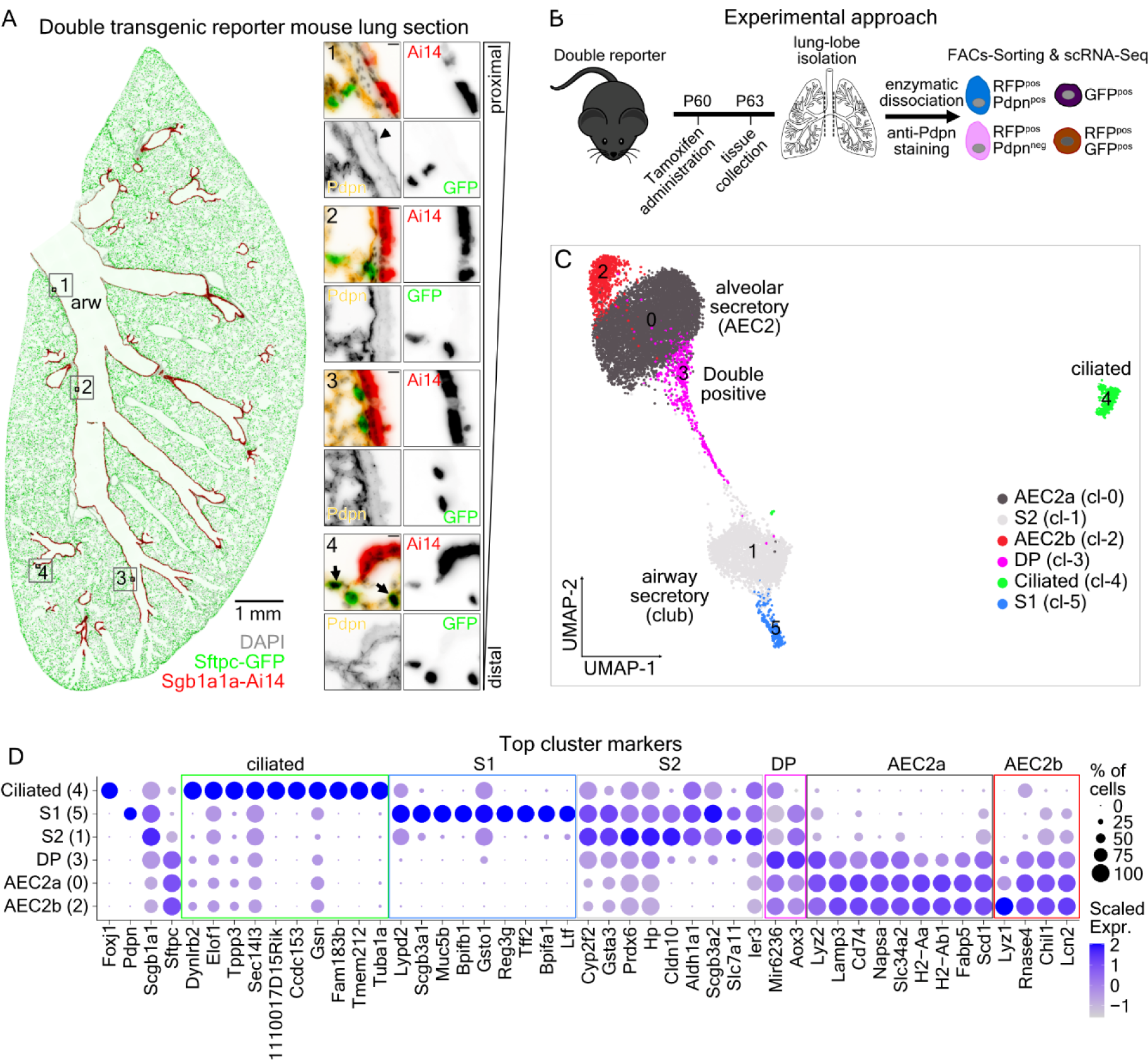
Characterization of lung secretory cell heterogeneity. **(A)** Colour-inverted fluorescence image of a left-lung section from an Scgb1a1-CreER^T2pos/neg^;Rosa26-Ai14^pos/neg^;Sftpc-GFP P63 reporter mouse, three days after Tamoxifen administration. DAPI (nuclei): grey, RFP (Ai14): red, GFP (Sftpc): green, Pdpn: yellow. Scale-bar: 1mm. Inserts correspond to the numbered ROIs. Insert Scale-bars: 20µm **(B)** Experimental outline for the isolation and single-cell RNA sequencing (scRNA-Seq) of the labelled cells from the reporter mouse. **(C)** UMAP-plot of 12030 cells, grouped into six clusters. Cluster (cl) 0: AEC2a (alveolar epithelial cell type-2a) (dark grey), cl-2: AEC2b (red), cl-5: airway secretory cell type-1 (S1) (blue), cl-1: airway secretory cell type-2 (S2) (light grey), cl-3: Scgb1a1^pos^ Sftpc^pos^ double positive (DP) (magenta) and cl-4: ciliated cells (green). **(D)** Balloon-plot of known epithelial markers. Ciliated: *Foxj1*, Club cells: *Scgb1a1*, Proximal-airway secretory and AEC1: *Pdpn*, AEC2 and DP-cells: *Sftpc*, in addition to the expression of the top-10 differentially expressed genes of each cluster. The genes were filtered according to average log2 Fold-change (>0.5), adjusted p-value (<0.001) and percent of positive cells (>0.25) and the top-10 markers according to average log2 Fold-change were plotted. Gene order follows the cluster order. Balloon size: percent of positive cells. Colour intensity: scaled expression (blue: high and grey: low).

We induced recombination in adult mice and three days later, we FACS-sorted the labelled cells for droplet-based scRNA-Seq (Fig. 1B, Extended Data Fig. 1A). We generated and analysed 12030 high-quality cDNA libraries (Extended Data Fig. 1B-C). The UMAP (Uniform Manifold Approximation and Projection) plot ^32^ and differential expression of marker genes in the clusters were consistent with the FACS-sorting criteria (Extended Data Fig. 1D-E). We annotated clusters according to the positivity for known lung epithelial cell markers (Fig. 1C-D, Extended Data Fig. 1F, Suppl. Table 1). A small cluster (cl-4) contained Foxj1^pos^ multiciliated cells found in the *Scgb1a1*creER-Ai14^pos^ Pdpn^neg^ cell sorting fraction (99.8%), suggesting that the modest *Scgb1a1* expression (Extended Data Fig. 1G) in ciliated cells was sufficient to induce recombination in few ciliated cells. As expected, cluster-4 cells uniquely expressed many genes related to cilium organization and function (GO:0044782) (Suppl. Table 2).

The *Scgb1a1*^pos^ airway secretory cells are separated into three clusters. Cluster-5 contains mainly Scgb1a1-Ai14^pos^ Pdpn^pos^ sorted cells that also expressed high levels of the proximal airway cell markers *Scgb3a1* and *Scgb3a2*^8^ (Fig. 1D). We annotated these cells as S1 (Secretory 1). Cluster 1 (cl-1) was almost exclusively (99.2%) composed of *Scgb1a1*creER-Ai14^pos^ Pdpn^neg^ cells (Extended Data Fig. 1D-E), which were also positive for *Kdr*^33^ (Extended Data Fig. 1F) and previously reported epithelial cell markers. We annotated these cells as S2. Cluster-3 contained 92.4% of Scgb1a1creER-Ai14^pos^ Sftpc-GFP^pos^ double-positive (DP) cells. The rest of these cells were evenly distributed among the two alveolar secretory (AEC2) clusters 0 and 2 (Extended Data Fig. 1D-E). This suggested that some AEC2s also express *Scgb1a1*. Cl-3 cells co-expressed moderate levels of S2 and AEC2 markers. However, we did not detect unique markers (Fig. 1C), suggesting that they represent an intermediate cell state between airway and alveolar secretory cells.

The clusters of alveolar secretory cells AEC2 (cl-0, AEC2a) and (cl-2, AEC2b) differed in *Lyz1* expression in the AEC2b cluster (Extended Data Fig. 1H), as reported previously^34^. Both alveolar clusters (AEC2a and b) expressed high levels of genes involved in lipid biosynthesis (GO:0008610) and lipid transport (GO:0006869), in addition to genes implicated in the regulation of leukocyte activation (GO:0002694), such as various MHC class-II genes involved in antigen presentation (Suppl. Table 2). Overall, this analysis defined two airway secretory cell types, a group of Scgb1a1^pos^ Sftpc^pos^ cells, two secretory alveolar cell identities that differ in the expression of *Lyz1* and a cluster of Foxj1^pos^ ciliated cells.

### Gene expression patterns suggest distinct secretory cell functions

The continuous arrangement of the lung secretory cells from the S1 to the AEC2b cluster in the UMAP-embedding suggested intermediate expression levels of cell-specific gene programs in the cells bridging the main bodies of each cluster. To explore the transcriptional heterogeneity along this continuum, we used diffusion maps^35^ and trajectory analysis (Fig. 2A), similar to the pseudotemporal cell ordering along a developmental trajectory. We ordered equal numbers of randomly selected cells from each cluster and identified 1563 differentially expressed genes (DEGs) along the trajectory. These genes can be grouped into 10 stable modules (Fig. 2B, Suppl. Table 3). The aggregated gene expression scores confirmed groups of co-expressed genes, which are either gradually reduced from S1- to S2-cells (modules-5 and-2) and gene programs graded in the opposite orientation (modules-4 and -1). Module-3 genes showed equal activation in both S1 and S2, but also in ciliated cells, representing a general airway gene expression program. Modules -7 and -8 were enriched in S2 cells, but only module-7 was expressed in ciliated cells. Module-9 genes were enriched in all but the S1 cells and modules-10 and -6 in all but S2 cells (Extended Data Fig. 1I). This analysis reveals shared, distinct and graded gene expression programs in the adult airways. Gene ontology (GO) analysis of the modules suggested selectively enriched biological processes for each cell type (Fig. 2C-D, Suppl. Table 3). For example, module-5 includes genes related to innate immunity regulation (GO:0002682, *Reg3g*^36^, *Ltf, Bpifb1*^37^ *P2rx4*^38^, *Il13ra1*^39^ and *Mfge8*^40^) and its expression is restricted to S1-cells only. Module-2 genes are also highly expressed in S1 cells but are gradually decreased in cells of the S2 and DP clusters. These genes encode various metabolic enzymes (e.g., *Gsta1*, *Acsl1* and *Gstm5*), cytokines (*Cxcl1*, *Cxcl2*, *Cxcl5* and *Cxcl17*) and interferon-induced antiviral proteins (*Ifitm1*, *Ifitm3* and *Ifit1*), suggesting differential functions of secretory cell-types in response to chemicals (GO:0070887) and viruses (GO:0034097). This is also supported by previous functional analyses of few module-2 genes. Transcription factor encoding genes (TF) *Six1*^41^ and *Spdef*^42^ have been implicated in airway inflammation and *Irf7* was upregulated in airway secretory cells upon RSV infection^43^.

**Figure 2.**
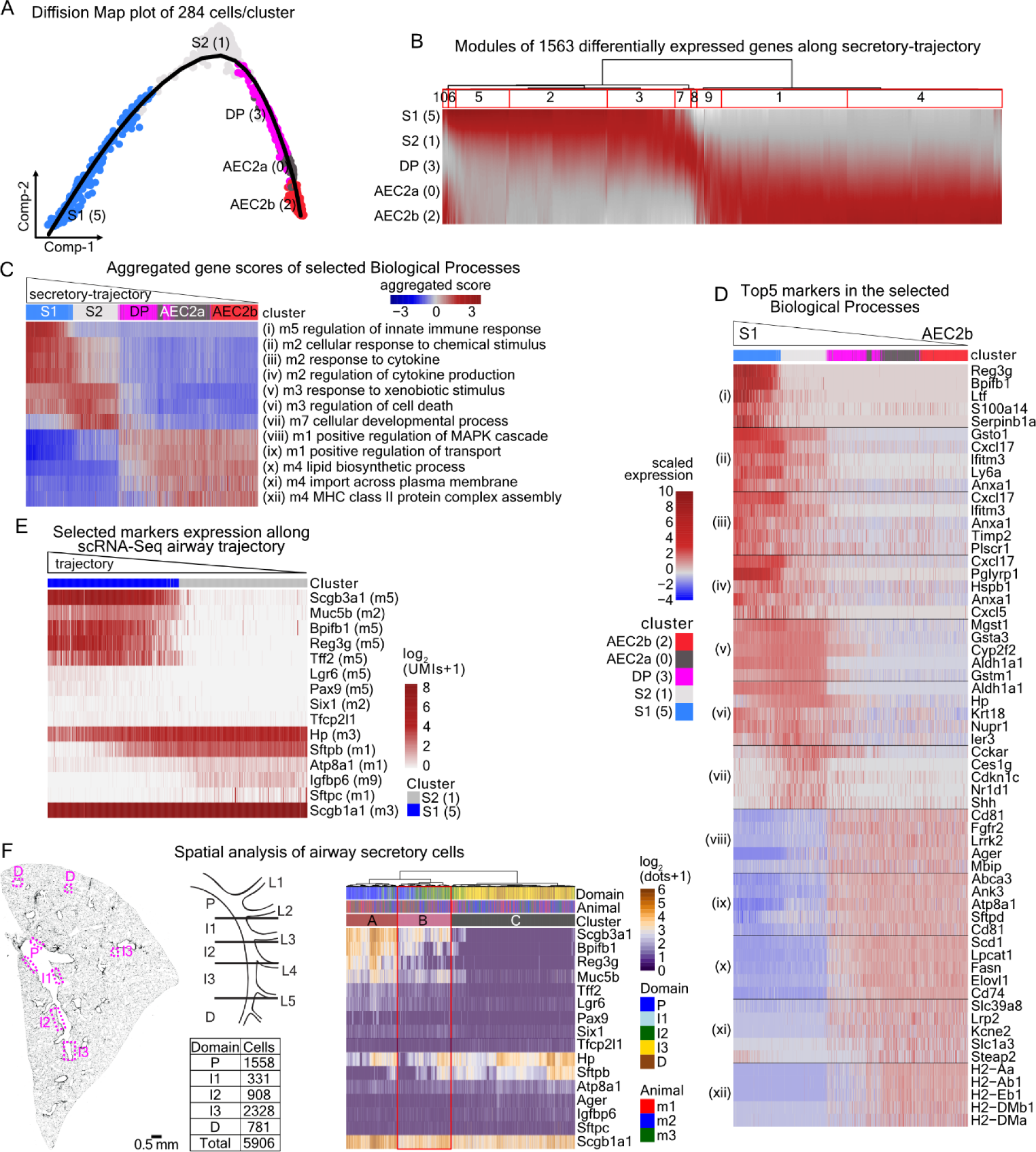
scRNA-Seq trajectory recapitulates the airway proximal-distal pattering. **(A)** Diffusion-map of secretory cell clusters. Colours and numbers as in Fig. 1C. Line: estimated pseudotime-trajectory by Slingshot. **(B)** Heatmap of the 1563 differentially expressed genes (FDR<0.001 and meanLogFC>1) along pseudotime, based on tradeSeq. The dendrogram of hierarchical clustering (left) indicates 10 stable gene-modules. Bootstrapping values: module-1: 0.65, module-2: 0.62, module-3: 0.61, module-4: 0.74, module-5: 0.68, module-6: 0.75, module-7: 0.8, module-8: 0.7, module-9: 0.64, module-10: 0.9. Colour intensity: scaled expression. Dark red: high, Gray: low. **(C)** Heatmap of the aggregated expression scores of the genes in the indicated biological processes (Suppl. Table3). The number after “m” indicates the module containing the genes in “B”. Cells were ordered according to pseudotime (Fig. 2A). red: high, blue: low. **(D)** Heatmap of the top-5 genes (according to “waldStat” score) of the indicated biological processes in “C”. Cells were ordered according to pseudotime. Colour: scaled expression (red: high, blue: low). **(E)** Heatmap of the selected S1 (cluster-5) and S2 (cluster-1) scRNA-Seq markers, ordered along pseudotime. Expression levels: log_2_(normalized UMI-counts+1) (library size was normalized to 10.000). **(F) Left panel:** (left) Representative adult mouse lung section stained with DAPI (grey) showing examples of imaged areas along the PD-axis (3 lungs have been analysed with similar results). (Right-up) Cartoon of airway domain classification approach. (Right-bottom) Synopsis of analysed cell-ROIs from three animals, for indicated domains. **Right panel:** Heatmap of 3096 analysed airway secretory cell-ROIs, showing the log_2_(SCRINSHOT dots +1) signal for the selected markers. Cell-ROI ordering is based on hierarchical clustering. Annotation bars show the (i) airway domains of the cell-ROIs, (ii) the analysed mouse and (iii) the indicated cluster. Cluster-A: P-domain cells 38.61 ± 6.89%, I1-domain cells 21.61 ± 2.46%. Cluster-B: I2-domain cells 37.79 ± 7.54%. Cluster-C: I3-domain cells 52.08 ± 6.57%, D-domain cells 69.49 ± 2.20%.

Module-3 genes are uniformly expressed in S1- and S2-cell clusters and primarily encode metabolic and detoxification enzymes (e.g., *Cyp2f2*, *Aldh1a1* and *Gsta3*), supporting the notion that a general function of airway secretory epithelium is to respond to xenobiotics (GO:0009410, GO:0006749) to detoxify inhaled air.

The cells in the S2-cluster selectively express module-7 genes, which relate to the general term cellular development (GO:0048869). This module contains developmental genes, such as *Shh*^44–46^ and the negative regulators of airway inflammation *Kdr*, *Sema3e*, *Sema3a* and *Nr1d1*^33, 47–49^.

Cells in the DP and in both AEC2 clusters show a selective and gradual upregulation of module-1 genes, which were decreased in the S2 and S1 clusters. Module-1 contains genes encoding kinases like *Fgfr2* and *Lrrk2*, required for maintenance and function of the alveolar type 2 cells^50–53^. Additionally, *Atp8a1*, *Abca3* and the Rab-family genes *Rab27a* and *Rab34* relate to phospholipid transport (GO:0051050) and vesicle trafficking.

Finally, the cells in the two alveolar epithelial clusters upregulate module-4 genes, which encompasses known regulators of alveolar cell differentiation and maintenance, like *Etv5*^54^ and *Nkx2-1*^55, 56^, together with genes related to lipid biosynthesis (GO:0008610) and transport across the plasma membrane (GO:0098739). Notably, the enriched expression of *H2-Eb1, H2-DMb1, H2-Dma, H2-Aa* and *H2-Ab1* relating to the MHC class-II complex assembly (GO:0002399) suggests a selective role of alveolar secretory cells in antigen-presentation to immune cells.

This analysis identifies several gene programs that are shared but also differentially expressed among the secretory cell clusters. The expression intensity of these programs along a continuum of cell states suggests that prominent biological processes of the lung epithelium relating to immune responses, detoxification, lipid biosynthesis and ion transport are segregated in the airway and alveolar compartments but are also expressed in a graded fashion along the secretory trajectory.

### Spatial analysis of the secretory cell trajectories in the airways

To further investigate the gene expression gradients along the cell trajectory between S1 and AEC2b cell-states we focused on the spatial analysis of marker genes in the airways. We first selected a panel of 18 DEGs (Fig. 2E) and detected their transcripts *in situ* by SCRINSHOT^57^. We quantified the signals in 5906 manually segmented airway epithelial cell-ROIs in distinct airway positions, based on the stereotyped airway branching pattern of the left lung-lobe^2^ (Fig. 2F, Extended Data Fig. 2A). Hierarchical clustering of the 3096 secretory cells showed high expression of module-5 genes, *Scgb3a1* and *Muc5b* in the proximal domains (P and I1), whereas module-1 genes, *Sftpb* and *Atp8a1* were more abundantly expressed in the distal domains I3 and D. Interestingly, cells in the intermediately located I2 domain co-expressed lower levels of both module-1 and module-5 markers (Fig. 2F), resembling the opposing graded expression of module 1 and module 5 genes along the secretory cell cluster trajectory (Fig. 2E). This suggests that secretory cell-states S1 and AEC2, expressing high levels of unique markers, are located at the proximal and distal sites of each airway branch and cells in intermediate branch positions are in an intermediate cell-state expressing moderate levels of both the S1 and AEC2 gene modules. To further test this hypothesis, we analysed protein expression levels by immunofluorescence co-staining for few S1 and S2 markers (Scgb3a1, Muc5b, Hp and Atp8a1) relative to E-cadherin, which is homogeneously expressed in the epithelium (Extended Data Fig. 2B-C). Also, this analysis revealed expression gradients, where S1 markers were highest in the P domain and gradually reduced in the I (1-3) and D domains. The relative levels of the S2 markers Hp and ATP8a1 formed an opposing gradient, highest in the D-domain and gradually reducing towards the P-domain.

Many of the module-5 and -2 genes relate to innate immune responses and they are predominantly expressed in proximal airway S1-cells of adult mice. We, therefore, examined if environmental factors or the lung microbiome sets their basal expression patterns. We selected antibodies against two S1 and two S2 markers to determine relative protein levels in germ-free and pathogen-free conditions relative to E-Cadherin. The expression levels of these markers showed similar distribution regardless of the different environmental exposures (Fig. 3A-D). This suggests that the localized gene expression programs of epithelial secretory cells are initially specified genetically. Several genes in these programs are known to be further activated upon infections, tissue damage or inflammatory disease^12^. The graded expression values in the spatial analysis (Fig. 2F) and in the scRNA-Seq diffusion maps (Fig. 2E) indicates that the trajectory of secretory cells reflects the PD pattern of gene expression in the airway secretory epithelium (Fig. 3E). The expression values may thus provide a conceptual ruler for positioning airway secretory epithelial differentiation states along each branch. The unaltered peaks and valleys of S1 and S2 marker expression in lungs of germ-free mice suggest that airway cell patterning can be influenced but is not initially dependent on microbes.

**Figure 3.**
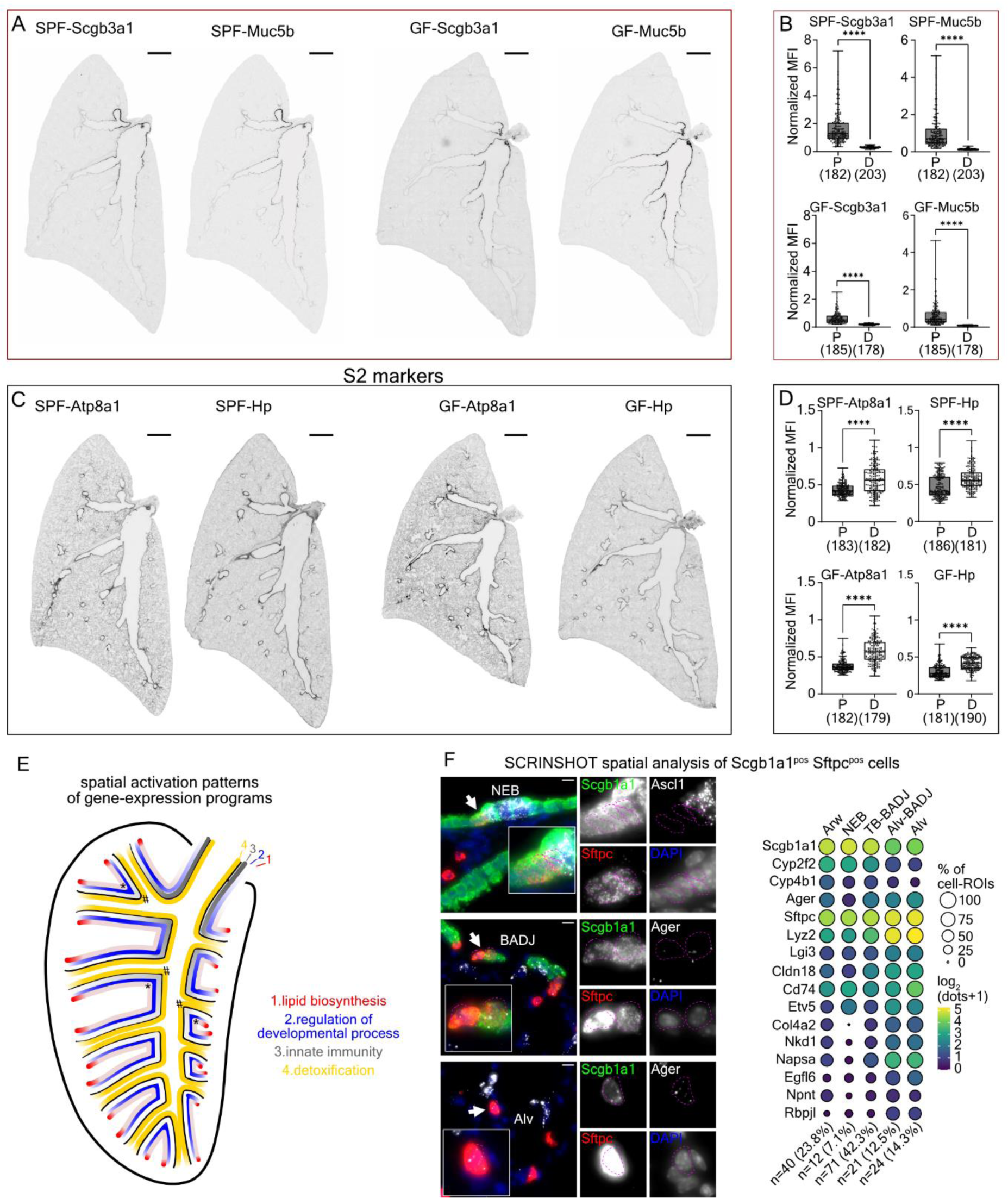
Gene expression patterns along the airway PD-axis. **(A)** Immunofluorescence stainings of whole lung sections of specific pathogen free (SPF) and germ-free (GF) 2 months-old mice for Scgb3a1 and Muc5b S1-markers. Scale-bars: 1000 µm. **(B)** Quantification of immunofluorescence mean fluorescence intensity (MFI) of the indicated target, normalized to the E-Cadherin signal. Numbers in parentheses: number of analysed proximal and distal cell-ROIs. Statistics with Student’s t-test: **** p<0.0001. **(C-D)** As in “A-B” for the Hp and Atp8a1 markers that are highly expressed in distal airways. **(E)** Graphical representations of the activated gene expression programs (as in Fig. 2C) along the proximal-distal axis of the adult mouse lung airways. Colour intensity: activation level. Dark: high, Fade: low. Exceptions in the expression of the lipid metabolism (asterisk) and detoxification (hash) programs relating to neuroepithelial body topology. **(F) Left:** SCRINSHOT analysis images for Scgb1a1^pos^ Sftpc^pos^ cells (arrows) close to neuroendocrine (NE) cells (upper panel), terminal bronchioles (TB) (middle panel) and alveoli (lower panel). Magenta dotted-lines: outlines of 2µm-expanded Scgb1a1^pos^ Sftpc^pos^ nuclei. Arrows: Scgb1a1^pos^ Sftpc^pos^ cells. Sftpc: red, Scgb1a1: green, Ascl1: grey and DAPI: blue. **Right:** Balloon plot of the 16 analysed genes (module-3: *Scgb1a1, Cyp2f2* and *Cyp4*b1, module-1: *Ager, Sftpc* and *Lyz2* and module-4: *Lgi3, Cldn18, Cd74, Etv5, Col4a2, Nkd1, Napsa, Egfl6, Npnt* and *Rbpjl*) in the 170 identified Scgb1a1^pos^ Sftpc^pos^ cells, according to their position. Balloon size: percentage of positive cells. The colour intensity: log_2_(SCRINSHOT dots +1). Yellow: high, Dark blue: low. “n”: number of cells in the specified position.

### Spatial analyses of the Scgb1a1^pos^ Sftpc^pos^ cells

We further examined topology-related gene expression markers in rare secretory cell identities like the double positive (DP) Scgb1a1^pos^ Sftpc^pos^ (DP) cells using an additional SCRINSHOT probe panel of 16 DEGs along the secretory trajectory, together with the neuroendocrine (NE) cell markers, *Ascl1* and *Calca*. We confirmed the previously reported localization of the DP-cells in bronchioalveolar-duct junctions (BADJs) and close to NEBs^58^ (Fig. 3F, Extended Data Fig. 2D). We found them mainly at airway terminal bronchioles (TB) (42.3%) and to a lesser extend in the alveolar compartment (12.5%) close to BADJs. A small fraction of DP cells (7.1%) was close to NE cells, within a 20 µm radius surrounding the neuroepithelial body (NEB) borders. We found that cell-ROIs in the TB-part of BADJs express higher levels of the S2- (module-3) than AEC2-enriched markers (module-1 and -4), whereas DP-cell-ROIs in the alveolar part of BADJs and alveoli showed an opposite profile. Immunofluorescence stainings for few protein markers confirmed the SCRINSHOT results (Extended Data Fig. 2E). We conclude that DP-cells are detected in three distinct locations along the epithelial PD-axis, expressing high levels of S2 or AEC2 markers, depending on their position. The DP-cells in the vicinity of NEBs may correspond to v-club cells^21, 22^ and were found more distantly from NEs than the *Upk3a^pos^* u-club cells^23^ (Extended Data Fig. 2F). We conclude that even rare secretory cell types express graded levels of S2 and AEC2 cell markers depending on their position in the airway tree. Their distribution and distinct identities may reflect unique signals from the NEB microenvironment, such as Notch-signalling^59^.

### In vitro differentiation potentials along the airway epithelial trajectory

The graded activation levels of distinct gene expression modules reflect a continuum of cell states along the airway with possible functional differences in the proximal, intermediate and distal regions. As a first test of this hypothesis, we isolated cells from double reporter mice (as in Extended Data Fig.1A) and cultured them in Matrigel together with cells from a mouse lung fibroblast cell-line (MLg-2908), as described before^60, 61^ to compare their cell proliferation and differentiation potential. The proximal-domain cells correspond to the sorted Scgb1a1creER-Ai14^pos^ Pdpn^pos^ fraction, the Scgb1a1creER-Ai14^pos^ GFP^pos^ (DP) to the airway distal end and the Scgb1a1creER-Ai14^pos^ Pdpn^neg^ cells derive from intermediate and distal airways. We initially assessed cell clonogenic potential and found that the two Scgb1a1creER-Ai14^pos^ GFP^neg^ cells fractions were more potent than DP-cells under our culture conditions (Fig. 4 A). Cultured cells produced three colony-types as previously described^60, 62^. The large cystic colonies (type-A: bronchiolar) expressed the airway secretory markers Scgb3a1, Muc5b, Hp and Scgb1a1 in addition to the ciliated cell marker acetylated tubulin^63^ and the basal cell marker Krt5^64^ and Pdpn. The dense colonies (type-C: alveolar) were positive for the alveolar markers Sftpc and Ager and the type-B (bronchioalveolar) colonies showed mixed morphology and expression of both bronchiolar and alveolar markers. The Pdpn^neg^ cells produced predominantly bronchioalveolar and alveolar colonies, in contrast to the mainly bronchiolar ones of Pdpn^pos^ and the exclusively alveolar ones of GFP^pos^ cells (Fig. 4B-E).

**Figure 4.**
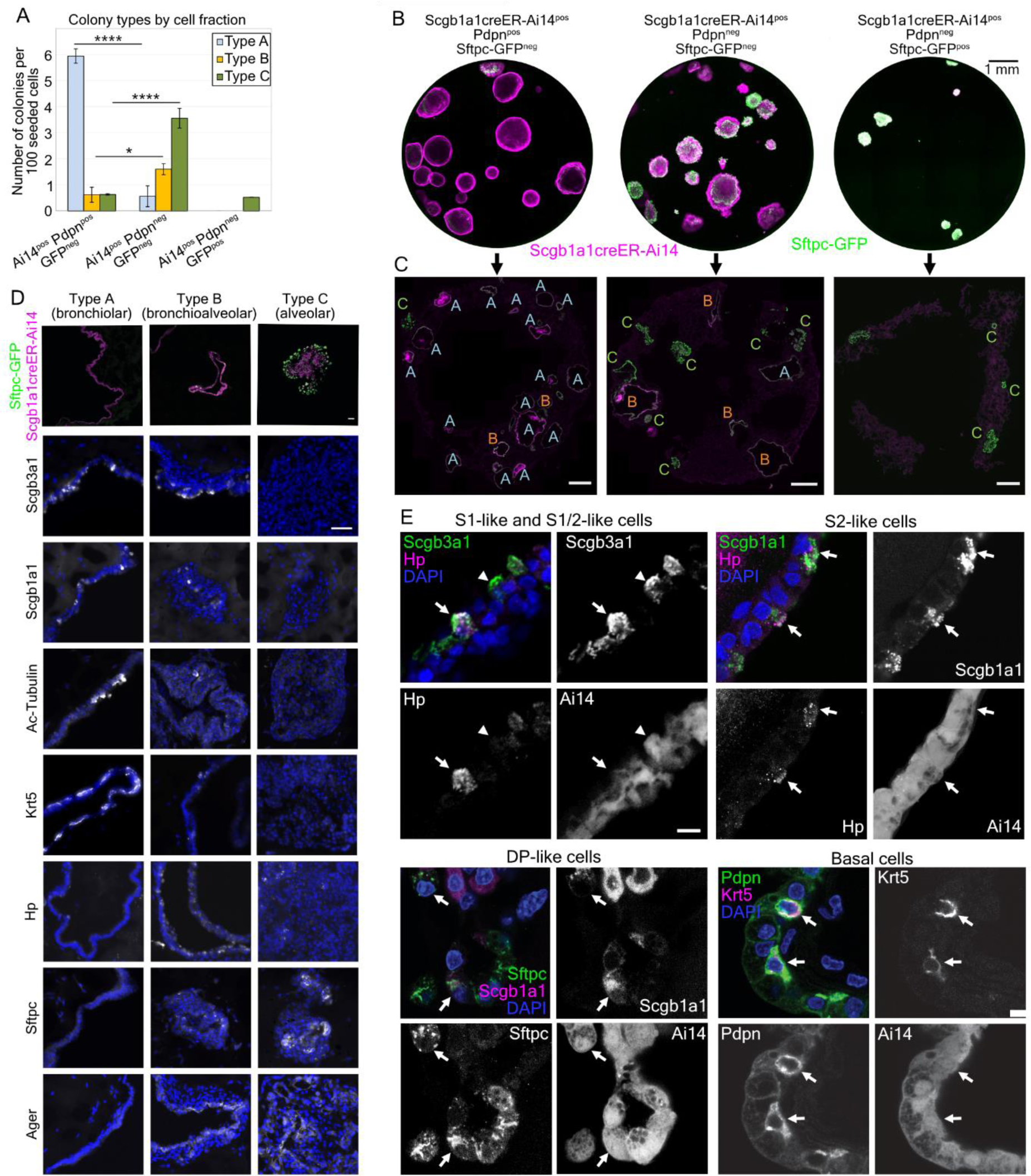
Clonogenic and differentiation potential of airway secretory cell states. **(A)** Bar plot of colony numbers per 100 seeded cells (counted 2-3 whole wells per cell type from four animals, three independent experiments) with embedded percentage of colony types derived from each type of seeded cells, (counted one well of each cell type from three animals, three independent experiments). Type-A: bronchiolar colony, Type-B: bronchioalveolar colony, Type-C: alveolar colony. Error bars correspond to standard error from the mean, * p<0.05, ** p<0.01, *** p<0.001. Number of colonies counted from 4 different mice, type-A: 52, type-B: 16, type-C: 44. Two tailed equal variance Student’s t-test (after a variance comparison test for all datasets) was performed. **(B)** Representative whole-culture images from Scgb1a1creER-Ai14^pos^ Pdpn^pos^ Sftpc-GFP^neg^, Scgb1a1creER-Ai14^pos^ Pdpn^neg^ Sftpc-GFP^neg^ and Scgb1a1creER-Ai14^pos^ Sftpc-GFP^pos^ seeded cells. Scale-bar: 1mm. **(C)** Images of 10µm thick sections of representative cultures showing Ai14 and GFP transgene fluorescence. The letters indicate the colony annotations. **(D)** Representative images of the three types of colonies, showing the Ai14 and GFP transgene fluorescence and the immunofluorescence signal for bronchiolar (Scgb1a1), S1 (Scgb3a1), basal (Krt5), ciliated (acetylated Tubulin), and distal epithelial (Hp, Sftpc, Ager) markers. Nuclei-DAPI: blue. Scale bar: 50 µm. **(E)** Confocal microscopy images for the detection of S1 (Scgb3a1^pos^ Hp^pos^), intermediate (Scgb3a1^pos^ Hp^pos^), S2 (Scgb1a1^pos^ Hp^pos^) and DP (Scgb1a1^pos^ Sftpc^pos^) cells in analysed colonies. Arrows indicate positive cells for both analysed markers. Scale-bar 10 µm. Whole well culture section from each fraction of cells was stained and analysed from three biological replicates.

Our results suggest that the airway secretory epithelial cells have different characteristics relating to their topology and that they retain them in vitro. The increased potency of Pdpn^neg^ cells to produce bronchioalveolar colonies with positive cells for all identified airway secretory, ciliated, basal and alveolar markers (Fig. 4A) suggests that S2 cells represent a heterogeneous cell population in intermediate and distal airways with higher plasticity, than those of S1 proximal airway cells and DP-cells, at least under the uniform *in vitro* co-culture conditions.

### Airway epithelial cells mature postnatally

To identify when the different airway cell states are specified and determine potential lineage relationships, we induced labelling of embryonic Scgb1a1^pos^ cells with a farnesylated GFP variant (Scgb1a1creER-fGFP)^19^ and lineage-traced them. We induced recombination at the onset of Scgb1a1 expression^19^, on embryonic day (E) 16 and FACS-sorted labelled progeny cells at E19.5 and postnatal days P2, P21 and P60 for full-length scRNA-Seq^65^ (Fig. 5A).

**Figure 5.**
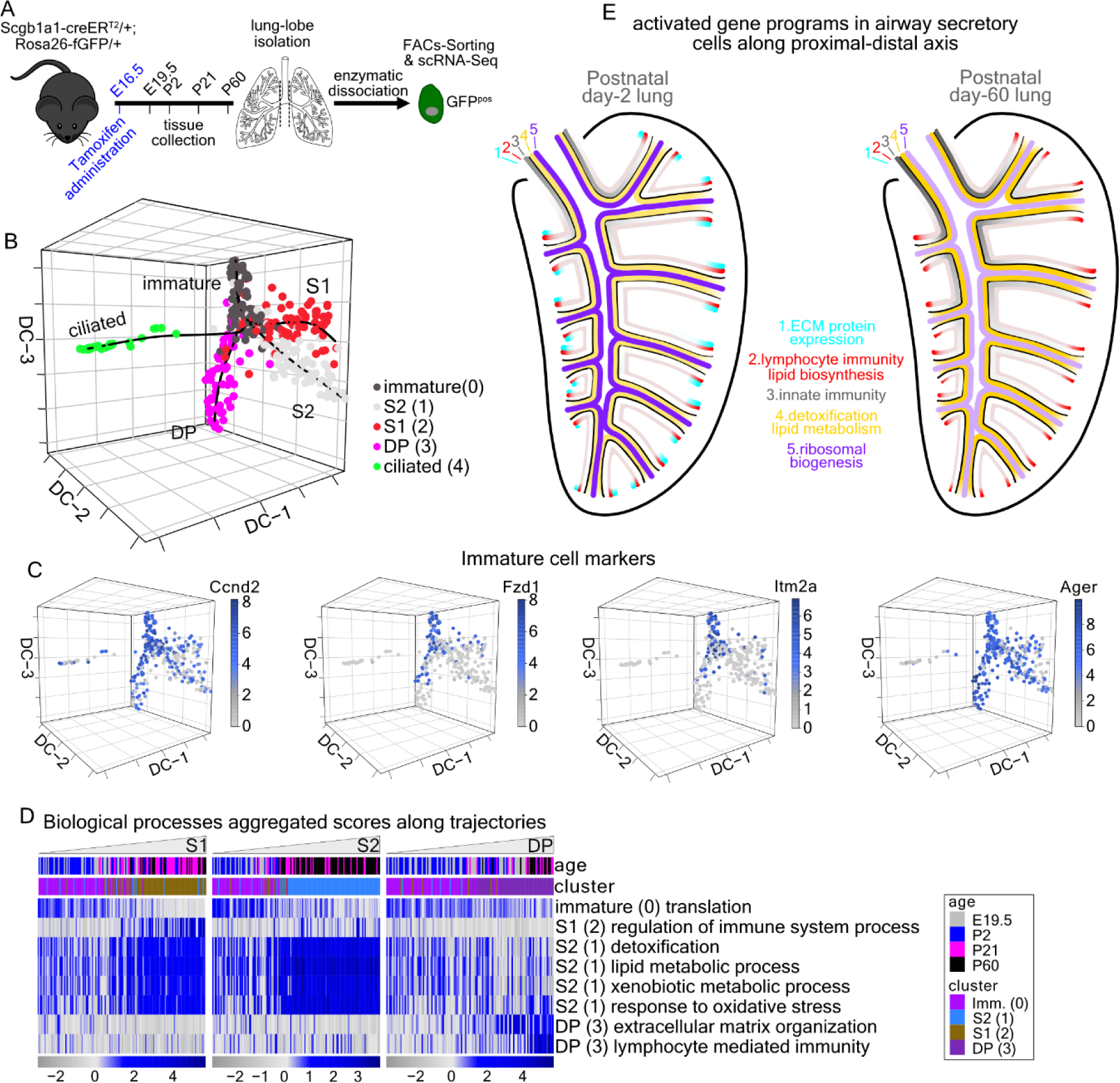
Lineage-tracing of airway secretory cell heterogeneity. **(A)** Experimental outline for the isolation and single-cell RNA sequencing (scRNA-Seq) of the labelled cells from the Scgb1a1-CreER^T2 pos/neg^;Rosa26-fGFP^pos/neg^ reporter mice. **(B)** 3D Diffusion-map plot of 354 full-length, single-cell cDNA libraries. Colours: suggested clusters. Lines: four distinct lineage-trajectories, calculated by Slingshot. **(C)** 3D Diffusion-map plots of the perinatally expressed genes *Ccnd2*, *Fzd1*, *Itm2a* and *Ager*. Expression levels: log_2_(normalized counts+1) (library size was normalized to 10^6^). Blue: high, Gray: zero. **(D)** Heatmaps of the aggregated gene expression scores of the indicated biological processes (see Suppl. Table 6). The cells were ordered according to the pseudotime values of the trajectories in “B”. Blue: high, Gray: low. **(E)** Synopsis of the gene expression programs activation in airway secretory epithelial cells, along the proximal-distal axis, in the postnatal day-2 (left) and -60 (right) lungs. Colour intensity: activation level. Dark: high, Fade: low. ECM: extracellular matrix.

We analysed 354 libraries (Extended Data Fig. 3A) using diffusion maps and trajectory analysis and found four distinct trajectories stemming from cl-0 composed of immature secretory cells from E19.5 and P2 lungs. The trajectories end in four mature clusters containing cells from P21 and P60 lungs (Fig. 5B, Extended Data Fig. 3B). According to DEG analysis and marker gene expression Cluster 3 (cl-3) corresponds to DP, (cl-1) to S2, (cl-2) to S1 and (cl-4) to ciliated cells (Extended Data Fig. 3C, Suppl. Table 5) in the adult-cell dataset. Differential expression analysis between the clusters revealed a high level of *Upk3a* and *Krt15*^36^ in the perinatal cluster. These cells also upregulate *Cccnd2* implicated in injury repair^66,67^, the WNT receptor *Fzd1*, the autophagy regulator *Itm2a*^68^ and the AEC marker *Ager*^69^. The enriched gene sets of the mature cell clusters largely overlap with those of the adult dataset (Fig. 5C, Extended Data Fig. 3C, Suppl. Table 5).

Next, we used GO-analysis to identify enriched biological processes in the clusters (Extended Data Fig. 3D and Suppl. Table 6) and scored the cells along the trajectories based on the expression of the corresponding genes. We found that the perinatal immature cells highly and transiently express ribosomal genes, indicating high levels of mRNA translation (GO:0006412) and ribosome biogenesis (Fig. 5D, Extended Data Fig. 3D, Suppl. Table 6). The mature airway secretory cell gene modules, encoding detoxification, oxidative stress responses, xenobiotic and lipid metabolism were gradually established along the S1 and S2 trajectories but not in DP-cells (Fig. 5D). This is represented by the expression levels of genes encoding representative enzymes, such as the *Aldh1a1*, *Fmo2* and *Gsta3* (Extended Data Fig. 3E). Interestingly, the representatives of the innate immunity term (GO:0002682) *Scgb3a1*, *Tff2* and *Muc5b* reached high expression only at the very end of the S1-trajectory (Extended Data Fig. 3F). This suggests either slower establishment of the S1 gene expression program or that mature cells from intermediate and distal domains were erroneously placed along that trajectory. To test this, we also analysed the cells according to the actual developmental age of their isolation and found that the P60 cells have generally higher expression levels of the markers than those isolated at P21 (Extended Data Fig. 3G). The cells along the DP-trajectory gradually increase their ability for lymphocyte-mediated immunity (GO:0002449), upregulating the *Cd74*, *Ctsc*, *Hc*, *Emp2*, *H2-Aa* and *H2-Ab1*. The middle part of the DP-trajectory contains perinatal cells that likely contribute to the local extracellular matrix (ECM) organization (GO:0030198), expressing high levels of genes like the *Col4a2*, *Spock2* and *Matn4* (Extended Data Fig. 3H).

In summary, we showed that different clusters of adult airway secretory cells derive from an embryonic secretory *Scgb1a1^pos^* population. Differentiating cells mature postnatally, acquiring their functional characteristics during the first three weeks after birth (Fig. 5E). Overall, the perinatal airway epithelium shows high ribosomal biogenesis, gradually decreasing over time. The gene programs of innate immunity (S1-trajectory) are established later than those involved in xenobiotic metabolism and reduction of reactive lipid aldehydes (S1- and S2-trajectories), suggesting that cell specification programs are activated sequentially. In the developing distal lung, the differentiating DP cells transiently contribute to ECM composition and gradually acquire the expression of antigen presentation genes, which are also expressed by the adult AEC2s.

### Fgfr2 promotes distal differentiation programs and restricts the proximal ones

Fibroblast growth factor (FGF) signalling is crucial for lung epithelial branching ^70^ and is later required for AEC2 differentiation and maintenance^50–52, 71–73^. Our gene expression analysis showed that *Fgfr2* is also expressed in the adult epithelial cells belonging to gene module-1, which shows high levels in AEC2 and DP-cells and gradually decreases in S2 and S1 airway cells (Suppl. Table 3, Extended Data Fig. 4A). We further detected *Fgfr2* expression in the perinatal airway secretory cells (Extended Data Fig. 4B) and differentially localized Fgfr2 protein by immunofluorescence (Extended Data Fig. 4C) in P2 lung sections. Co-stainings with a Fgfr2β(IIIb)-Fc chimeric protein to detect the spatial distribution of Fgfr2-ligands together with Fgfr2 showed a punctate staining for the ligand, which was higher around the TBs and distal airway epithelial cells and lower at proximal airways. This suggested a more robust pathway activation in the distal airway regions^73^.

To examine if Fgfr2 signalling has any role in the postnatal establishment of gene expression gradients along the airway epithelium, we deactivated the receptor just after birth. We induced tamoxifen-mediated Fgfr2-inactivation^74^ in the *Scgb1a1* cells (Scgb1a1creER-Fgfr2KO) and utilized Rosa-loxTomato expression^30^ to detect recombination, and presumed mutant cells. After three Tamoxifen injections (P1-P3), we analysed the lungs at P7 by scRNA-Seq and histology (Fig. 6A). We clustered and annotated 9911 scRNA-Seq libraries from Epcam^pos^ cells from wildtype (library-1), Epcam^pos^ from mutant (library-2) lungs and Epcam^pos^ RFP^pos^ cells from mutant lungs (library-3). The UMAP-plot was consistent with the FACs-sorting criteria and showed that the inactivation did not affect *Fgfr2* expression in basal and AEC2 cells (Extended Data Fig. 4D-G). The alveolar (cl-0, -1, -5), basal (cl-7), NE (cl-8), ciliated (cl-6) and S1 (cl-4) clusters were composed of intermingled wildtype and mutant cells, indicating that there is no significant effect of the *Fgfr2* inactivation on these cells. In cluster-2, composed of S2 cells, RFP positive and negative cells showed a conspicuous separation, but we also detected *Fgfr2* transcripts in the cells from the *RFP*^pos^ libraries (Extended Data Fig. 4E), suggesting escaper cells, which recombined the Rosa26R-Ai14 (RFP) but failed to deactivate both *Fgfr2* alleles. This partial Fgfr2 inactivation was also validated by antibody stainings (Extended Data Fig. 4H) and led us to filter out the S1- and S2-cells with *Fgfr2* transcripts from library-3 before further analyses to reduce noise.

**Figure 6.**
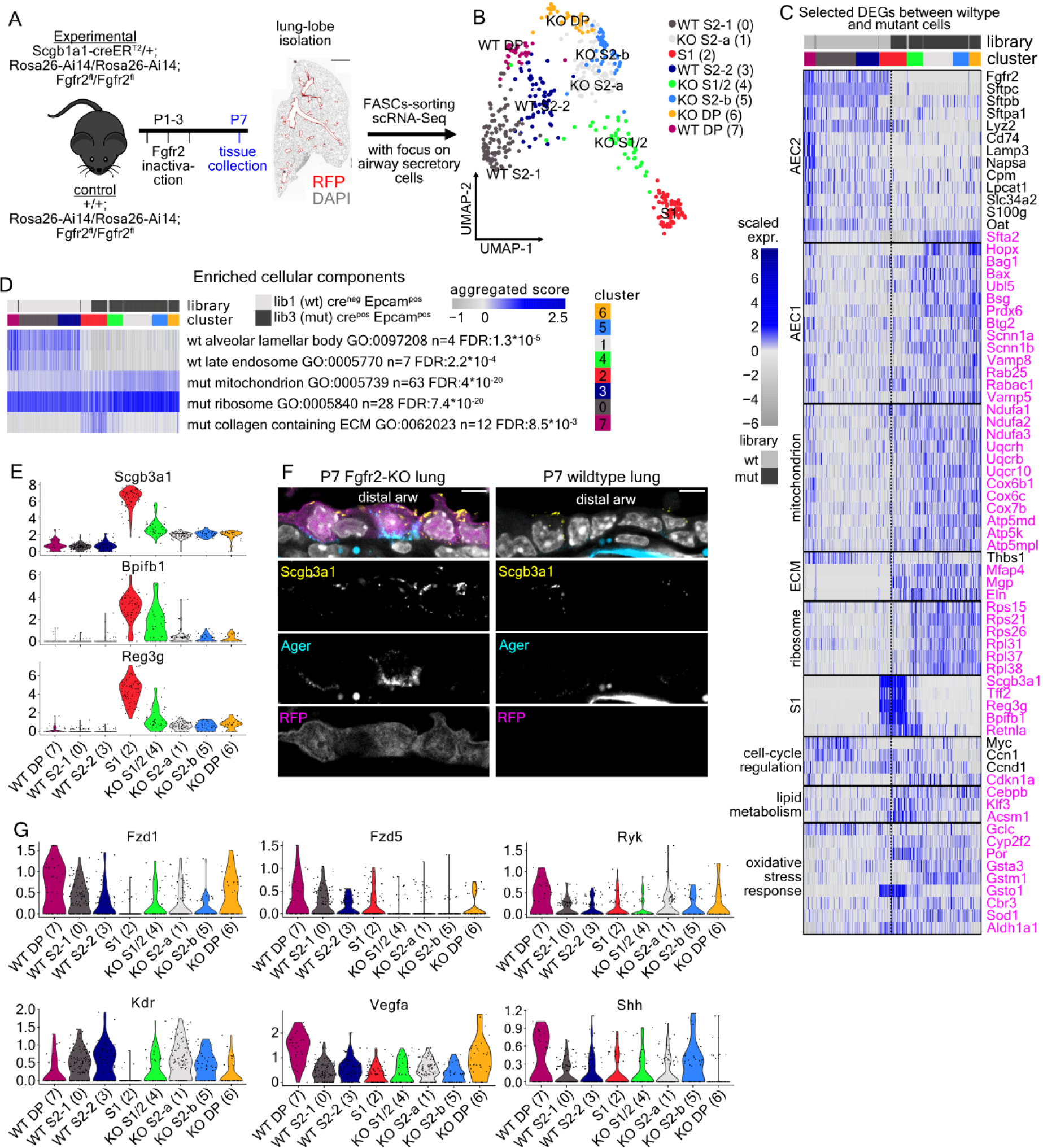
Fgfr2-inactivation in airway secretory cells causes extended gene expression changes. **(A)** Experimental outline for the perinatal inactivation of Fgfr2 in Scgb1a1^pos^ cells and analysis with single-cell RNA sequencing (scRNA-Seq) and histology. **(B)** UMAP-map plot of equal numbers of randomly-selected mutant and wildtype airway secretory cells of clusters -2 and -4 in Extended Data Fig. 4F. Colours: suggested clusters. **(C)** Heatmap showing the expression of selected, differentially expressed genes between the wildtype (library-1) and the mutant (library-3) airway secretory cells. Genes are organized in distinct categories according to previous knowledge and Gene Ontology analysis. Colour: scaled expression. blue: high, grey: low. **(D)** Heatmap of the aggregated scores of selected statistically significant, altered cellular components according to GO-analysis (see Suppl. Table 8). The results are based on the statistically-significant, differentially expressed genes between the wildtype (library-1) and the mutant (library-3) airway secretory cells. The cells are ordered according to the clusters (colours as in “B”). Score: blue: high, grey: low. “FDR”: false discovery rate, “n”: number of genes. **(E)** Violin plots of the *Scgb3a1*, *Bpifb1* and *Reg3g* showing their up-regulation in Fgfr2-mutant cells **(F)** Confocal microscopy, single-step images of distal airway epithelium from a postnatal day-7 (P7) Fgfr2-mutant lung (left) and a wildtype littermate (right). Immunofluorescence for Scgb3a1 (Yellow) and Ager (Cyan). Rosa26-Ai14 (magenta) indicates cells that underwent recombination. Nuclei-DAPI: grey. Scale-bar 5 µm. “arw”: airway. Three lungs for each condition were analysed. **(G)** Violin plots of the genes encoding Wnt receptors *Fzd1*, *Fzd5* and *Ryk*, the *Kdr* and its ligand *Vegfa* and *Shh*. In all violin plots, expression levels: log_2_(normalized UMI-counts+1) (library size was normalized to 10.000). Colours as in “B”.

We clustered equal airway secretory cell numbers from wildtype (lib-1) and mutant (lib-3) libraries and found eight clusters. These correspond to a single S1-cluster composed of wildtype and mutant cells, two wildtype S2- (WT S2-1 & WT S2-2) and three mutant S2-clusters (KOS2-a & KO S2-b), of which one expresses elevated levels of S1 markers (KO S1/2). In addition, we identified wildtype and mutant DP cell clusters (WT DP & KO DP) according to their DEGs (Fig. 6B, Extended Data Fig. 5A, Suppl. Table 7B-C). There was no clear pairwise correlation of the S2 and DP between the wildtype and mutant clusters (Extended Data Fig. 5B-C). To avoid comparing irrelevant cells, we compared all wildtype to all mutant airway secretory cells, regardless of clustering and identified 240 statistically significant DEGs, which we categorized based on GO-analysis and previous knowledge (Fig. 6C-D, Suppl. Table 7D-F and 8). We also related the expression levels of the affected genes in the *Fgfr2* inactivation experiment with their levels in the lineage-tracing experiment (Extended Data Fig. 5D). We included all previously defined epithelial cell types regardless of genotype (Extended Data Fig. 5E) to identify both possible general, temporal and cell-type specific differentiation defects.

We found a prominent reduction of AEC2 marker levels (*Sftpc*, *Napsa*, *Cd74)* accompanied by increased expression of the AEC1 TF *Hopx* ^75^ (Fig. 6C). Mutant cells also upregulated a large number of nuclear-encoded mitochondrial genes encoding complex I, III, IV and V components, sodium channel genes (*Commd3*, *Scnn1a*, *Scnn1b*^76^) and genes relating to autophagy and vesicle trafficking (*Creg1*^77^, *Vamp8*, *Vamp5*^78^, *Rab25* and *Rabac*), which are all normally expressed by P42 AEC1s^79^ (Fig.6C, Extended Data Fig. 5E and 6A-C). The senescence- and cell-survival-related genes *Cdkn1a* (p21)^80, 81^, *Bax* and *Bag*1^82^ are also enriched in AEC1s and become up-regulated in mutant secretory cells. Interestingly, *Hopx*, *Vamp5, Creg1* and *Scnn1b* are also detected at low levels in adult wild-type S2 cells, indicating their propensity to further activate AEC1 programs upon signalling (Extended Data Fig. 6D). This suggests that Fgfr2 activation in distal airway cells up-regulates AEC2-related genes that are responsible for surfactant biosynthesis and lamellar body formation (Fig. 6D) and directly or indirectly down-regulates numerous AEC1 genes, that relate to mitochondrial function, ion homeostasis, vesicle trafficking and cell-survival. Fgfr2-inactivation also altered the expression of several genes involved in lipid metabolism, trafficking and adipogenesis (Suppl. Table 7F).

Fgfr2-inactivation in the airways also caused increased expression of ECM protein-encoding genes (*Eln*, *Mgp* and *Mfap4*) which are normally transiently expressed along the DP-cell trajectory (Extended Data Fig. 5D) and in developing AEC1 and AEC2 (Extended Data Fig. 6C). Similarly, mutant cells failed to downregulate numerous ribosomal-subunit genes, which are highly expressed in all immature lung epithelial cells and gradually decrease as specification proceeds (Extended Data Fig. 5D, 6C). These findings suggest that Fgfr2 is required for the normal progression of differentiation in distal airway epithelial cells.

The most significantly reduced TF in mutant S2-cells is *Myc*, which is normally highly expressed in immature airway secretory cells and becomes downregulated in mature S1 and S2 (Extended Data Fig. 5D, Suppl. Table 7G). Other significantly changed TFs are the down-regulated *Atf4* and *Ets1* and the up-regulated *Cebpb*.

An intriguing phenotype of Fgfr2-inactivation is the appearance of a new cell cluster of S2-cells (cl-4, S1/S2), which expressed increased levels of S1 innate immunity marker genes, such as *Tff2*, *Bpifb, Reg3g* and the *Scgb3a1* proximal cell marker (Fig. 6C, E). This suggested that Fgfr2 activation in distal cells restricts the S1-related gene expression program. We confirmed this observation by antibody stainings in mutant lungs, where Scgb3a1 protein was detected in distal airway epithelial cells co-expressing Ager (Fig. 6F). Similarly, Muc5b expansion was observed in adult Fgfr2-mutant distal airway secretory cells upon naphthalene-induced injury^83^.

To elucidate potential cell-autonomous or indirect mechanisms involved in this restrictive function of Fgfr2-signaling, we interrogated the transcriptomes of mutant and wildtype cells for the expression of genes encoding *Vegfa*, its receptor *Kdr* and *Ryk*, a Wnt co-receptor. These three genes are normally expressed in secretory cells and are required to restrict the proximal gene expression programs and mucus metaplasia upon airway epithelial injury^33, 84^. We found that the levels of both *Vegfa* and *Ryk* were reduced in the *Fgfr2* mutant DP- and S2-cells (Fig. 6G), suggesting that Fgfr2 activation in distal cells may also restrict the expression of proximal genes by activating the expression of relaying signals and receptors. We also observed a decrease in the levels of Shh^72^ in *Fgfr2* mutant DP cells (Fig. 6G), suggesting altered paracrine signalling from the distal epithelial cells to endothelial and other mesenchymal cell types of the mutant lungs.

In summary, perinatal Fgfr2-signalling in airway secretory epithelium promotes progression towards differentiation, specifies the levels of S2 and DP cell programs and at least indirectly changes airway patterning by activating genes encoding signals and receptors.

## Discussion

Our large-scale scRNAseq and spatial analysis of airway secretory cells in the adult lung suggests that cell characteristics are defined by a uniformly activated gene program relating to cell responses to xenobiotics and two opposing and partially overlapping graded programs. These programs encode genes relating to innate immunity, cytokine production and response to cytokines in proximal regions and lipid synthesis, surfactant production and antigen presentation in the distal ones (Fig. 7A). Similarly, recent reports showed graded expression patterns of a few distal markers in the distal human airway epithelium^7, 85, 86^. Future experiments are needed to investigate the presence of opposing gradients along the airway network in human donor samples to establish if the mouse patterns are conserved. Why may these developmentally controlled gene expression gradients be relevant for airway structure and function? Firstly, their slopes correlate with the tapering of airway branches, suggesting that graded gene expression programs may control branch size and shape, facilitating seamless airflow to the alveolar compartments. Second, the differentially localized expression of different types of immunity programs suggests that proximal cells are better endowed to present immediate innate responses and cytokine signalling. In contrast, the distal ones are more specialized for antigen presentation. The compartmentalization of immune functions correlates with the higher expression of mucin coding genes and greater abundance of multiciliated cells in proximal regions, where pathogens become trapped, targeted by antimicrobial peptides and propelled out of the tubes. Escaping pathogens may be further detected by distal airway cells, internalized and presented to lymphocytes, activating slower but long-lasting immune responses. Third, the gradients may reflect developmentally controlled positioning of cells expressing lower levels of specification genes in intermediate positions of each branch in the airway network. Such cells with higher plasticity and increased differentiation potential may efficiently and rapidly repair local damage caused by pathogens and inhaled toxic substances. Our *in vitro* culture experiment, comparing the differentiation of S1, S2 and DP cells, supports this notion, but our transcriptome analysis failed to define any specific markers genes for such cells, precluding their labelling and isolation at present.

**Figure 7.**
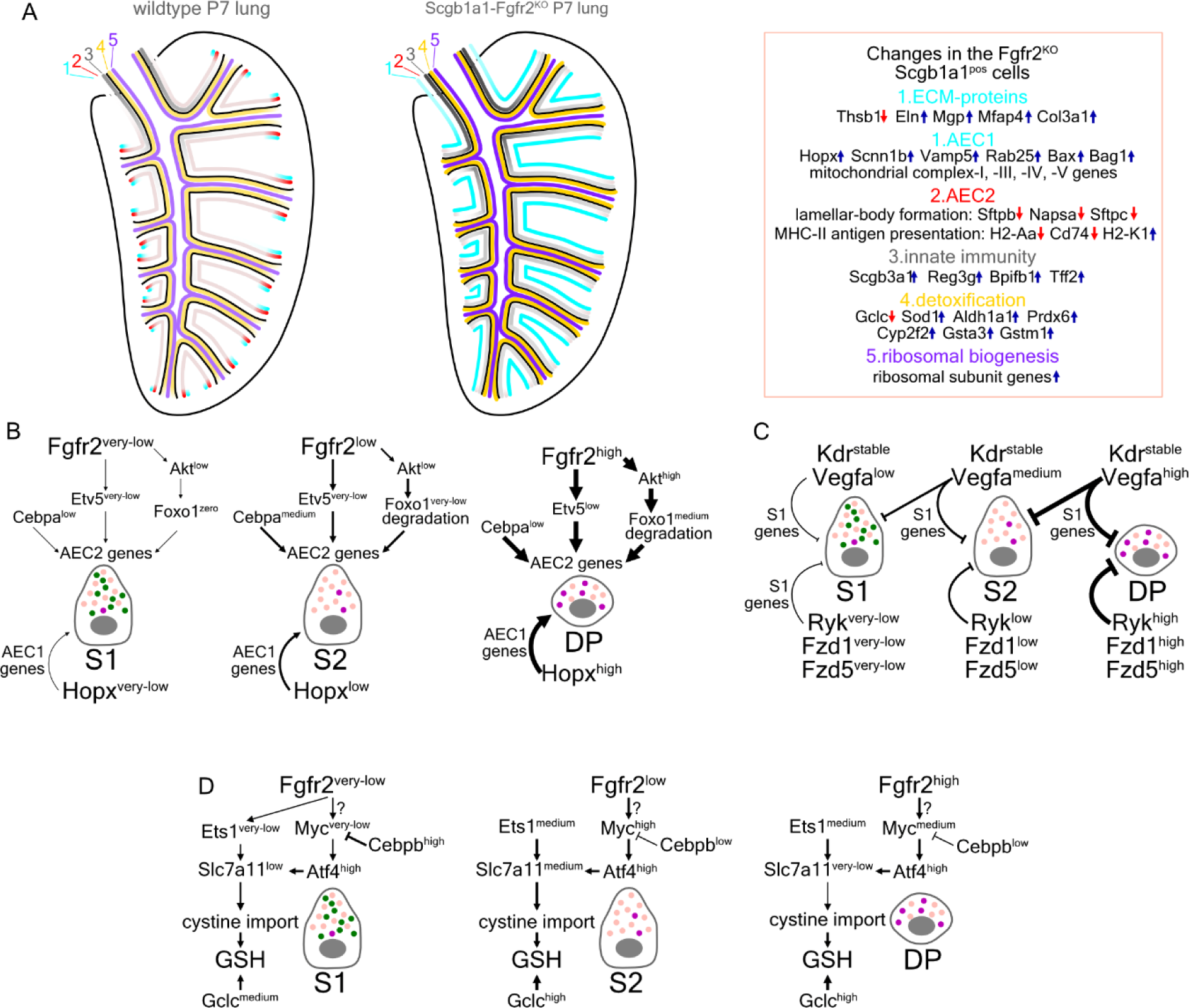
Synopsis of the Fgfr2-inactivation effects on perinatal airway secretory epithelium. **(A)** Schematic representation of the airway epithelium in postnatal day-7 wildtype (left) and Scgb1a1-Fgfr2^KO^ (right) lung. Colour intensity: activation level. Dark: high, Fade: low. ECM: extracellular matrix, AEC2: alveolar secretory cell type-2 genes, AEC1: alveolar secretory cell type-1 genes. **(B)** Model for the role of Fgf-signaling and Hopx in the regulation of AEC2 and AEC1 genes in the secretory cell populations along the airway epithelium. **(C)** Model for the role of Vegfa/Kdr pathway and Wnt-signaling in the regulation of S1-related gene expression programs in the secretory cell populations along the airway epithelium. **(D)** Model for the role of Fgf-signaling in the expression of genes involved in production of glutathione (GSH) in the airway secretory cells.

The lineage tracing analysis of the airway secretory cells and the meta-analysis of the GSE149563^79^ dataset enriched for alveolar-epithelial cells (Extended Data Fig. 5D, 6C) indicate that immature secretory cells downregulate the high expression of genes encoding ribosomal proteins as they become specified postnatally. Similarly immature secretory cells downregulate genes coding for ECM proteins as they reach the end of their differentiation trajectory. The expression of these genes is retained longer in DP-cells, where it becomes downregulated later. Immature secretory cells first upregulate a detoxification-related genetic program, which is retained and increased in S1- and S2-cells but becomes repressed in DP cells. Innate immunity-, lipid metabolism- and lymphocyte-mediated immunity gene programs become selectively established later in S1- and S2-cells. These results reveal common and distinct cellular mechanisms of secretory cell-type differentiation. Their sequential emergence suggests that they are hierarchically coupled.

The scRNAseq analysis of conditional Fgfr2-inactivation in the postnatal airway secretory epithelium (Fig. 7A) suggests a central role for distal Fgfr2-signalling in differentiation progression and airway patterning. First, the graded distal programs encoding surfactant production and endosomal vesicle traffic, which are normally expressed in DP and S2 cells, become severely reduced. Instead, DP- and distal S2-cells upregulate AEC1s gene expression programs, including genes encoding mitochondrial proteins and autophagy (Fig. 7B, Suppl. Fig. 1). This phenotype is similar to the one generated by the perinatal Fgfr2-inactivation in AEC2s, which reprograms them to AEC1s^51^. Second, our analysis reveals a long-lasting role of Fgfr2-signalling in all S2-cells, because the timed repression of ribosomal and ECM gene expression remains active in all S2-cells. This suggests that Fgfr2-signaling promotes the progression of secretory cell differentiation. Third, mutant S2-cells in intermediate positions along the airway network activate the S1, innate immunity-related gene program. This shorter-range effect may be mediated by additional, relaying signalling mechanisms involving Vegfa and Wnt-signaling (Fig. 7C, Suppl. Fig. 1).

A potential model on how Fgfr2-signaling may control detoxification in secretory cells derives from the severe downregulation of *Myc, Atf4* and upregulation of *Cebpb.* In previous studies, Myc induces *Atf4* expression in cancer cells and they co-operatively regulate promoters of various target genes, including *Cebpb*^87^. In return, Cebpb directly binds to the *Myc* promoter and inhibits its expression^88^. The reduced expression of both *Atf4* and *Ets1* in S2 mutant cells may be linked to lower *Slc7a11* expression, affecting the detoxification ability of the cells by compromising their ability to exchange intracellular glutamate with extracellular cystine for glutathione synthesis^89^. In our data, both *Slc7a11* and *Gclc* were reduced in mutant cells. Gclc is a rate-limiting enzyme of the first step of glutathione biosynthesis^90^. Reduced glutathione levels might, in turn, indirectly up-regulate other detoxification- and oxidative stress-related genes, like *Sod1, Aldha1*, *Gsta3* and *Gstm1,* in a presumed compensatory mechanism to spare the mutant cells from reactive oxygen species and lipid peroxidation (Fig. 7D, Suppl. Fig. 1).

Some developmental gene expression changes in the airways of Fgfr2 mutants show striking similarities with prior descriptions of cellular pathologies such as mucus hyperplasia rising during lung inflammation^33^ and small airway proximalization in smokers and COPD patients^91,92^. Our systematic description of the spatial organization of airway gene expression programs, their timely establishment and their regulation by Fgf-signalling along the mouse airways may help further molecular understanding of lung inflammation and COPD pathogenesis and define new avenues for treatments.

## Materials and Methods

### Animal models and Tamoxifen administration

All mouse experiments were performed according to Swedish animal welfare legislation and German federal ethical guidelines. The Northern Stockholm Animal Ethics Committee approved the project (Ethical Permit numbers N254/2014 and 15196-2018). The Research Animal Ethics Committee in Gothenburg approved the analyses of germ-free (GF) mice (Ethical Permit number 4805-23). The GF mice were maintained in flexible film isolators (Class Biologically Clean, Madison, WI, USA). GF status was monitored regularly by aerobic and anaerobic culturing and PCR for bacterial 16S rRNA. All mice were group housed in a controlled environment (room temperature of 22 ± 2 °C, 12h daylight cycle, lights off at 7 pm), with free access to autoclaved chow diet (#T.2019S; Envigo) and water. Breedings and experiments performed in JLU, Giessen, Germany were under the Ethical Permit with number GI 20/10, Nr. G 21/2017. For the lineage-tracing experiments, we used Scgb1a1-CreER^T2 pos/neg^;Rosa26-fGFP^pos/neg^ ^19^ mice. Noon of the day of the vaginal plug was considered as embryonic day (E) 0.5. We induced recombination by one oral dose (gavage) of Tamoxifen solution in corn oil (30mg/kg body weight) on E16.5, as described previously^19^. For the analysis of adult-lung epithelial heterogeneity and organoid cultures, we used Scgb1a1-CreER^het^;Rosa26-Ai14^het^;Sftpc-fGFP^het^ adult mice^29, 30^ and administered one Tamoxifen dose (100mg/kg body weight), 72 hours prior tissue collection. Experiments for Fgfr2-inactivation were performed using Scgb1a1-CreER^T2 pos/neg^;RosaAi14^pos/pos^;Fgfr2b^fl/fl^ and Scgb1a1-CreER^T2 neg/neg^;RosaAi14^pos/pos^;Fgfr2b^fl/fl^ mice. Tamoxifen was injected subcutaneously (87 mg/ kg body weight) on P1, P2 and P3 to induce efficient recombination.

### Tissue collection

Animals were euthanized by an intraperitoneal injection of anesthesia overdose, followed by incision of the abdominal vein. For embryonic lungs, we did not perform heart perfusion. For postnatal lungs, the chest was opened and the left atrium was excised. Lungs were perfused through the right ventricle of the heart with ice-cold PBS 1X pH7.4, using a 26G needle and 5ml syringe until they became white. Lungs were inflated with a mixture of 4% PFA:OCT (2:1 v/v) using a 20G catheter (Braun, 4251130-01), until the accessory lobe was expanded. The trachea was ligated (with silk 5/0 Vömel thread, 14739) and tissues were later fixed. For histological analysis, tissues were collected on E19.5, on post-natal day 2 (P2), 5 (P5), 7 (P7), 21 (P21) and 60 (P60). Embryonic and P2 tissues were fixed with freshly prepared 4% Paraformaldehyde (Merck, 104005) solution in PBS 1X pH7.4 (Ambion, AM9625) for 4 hours. Later stages were fixed for 8hours. Thereafter, the tissues were placed in OCT: 30% sucrose in PBS (2:1 v/v) over-night (O/N) at 4°C with gentle shaking and frozen in OCT (Leica Surgipath, FSC22), using plastic molds (Leica Surgipath, 3803025), by placing them in isopentane and dry ice. Tissue-OCT blocks were kept at −80°C until sectioning.

### Tissue dissociation and cell isolation

For full-length (Smart-Seq2) library preparation^65^, the left lungs were used for enzymatic digestion and the right lungs were treated as described above for histological analysis. For cell culture and droplet-based scRNA-Seq, both lungs were processed for digestion. Briefly, we cut the lungs in small pieces using a razor blade and digested them with elastase (Worthington, LS002292) and DNase-I 0.5mg/ml (Sigma-Aldrich, DN25) in HBSS (Gibco, 14175), at 37°C for 1 hour with rotation. An equal volume of HBSS++ [HBSS (Gibco, 14175), supplemented with 2% 0.2µm filtered FCS (Gibco, 10500), 0.1M HEPES (Sigma-Aldrich, H0887), antibiotics (Gibco, 15240096) and EGTA 2mM was added and the suspension was mixed gently. Then, the cells were centrifuged at 800g for 10 minutes at 4°C. The supernatant was removed with a serological pipette and cells were resuspended in HBSS++. Viability was tested using trypan blue (Sigma-Aldrich, T8154) (1:1 dilution) and the presence of fluorescence-positive cells was evaluated using a fluorescence microscope. Before sorting, cells were passed through a 100µm BD Falcon (BD Biosciences, 340610) to remove cell aggregates.

For E16.5 lungs, after digestion, centrifugation and resuspension in HBSS++, cells were passed through a 100 µm filter (BD Biosciences, 340610) and counted. We resuspended them in HBSS++ to obtain 20×10^6^ cells/ml. 100µl of cell suspension were stained with 0.5µl of anti-EpCam-PE antibody (Biolegend, 118205) and 0.5µl anti-CD45-APC antibody (Biolegend, 103112). Replicates of the above reactions were set in separate tubes to prevent aggregate formation, which is typical when a large number of epithelial cells are centrifuged.

For cell-sorting, we used a BD FACSARIA III cell-sorter with 100µm nozzle using single-cell sorting purity. Cells from all stages were isolated according to GFP and/or Tomato (for Rosa-Ai14 mice) expression. Non-transgenic and single-transgene (either Scgb1a1-CreER;Rosa-Ai14 or Sftpc-GFP) positive animals were used for instrument calibration. For cell culture experiments, cells were sorted in HBSS++ medium and for droplet-based sequencing, we omitted EGTA and HEPES, according to 10x Genomics instructions.

### scRNA-Seq of adult lung cells

Droplet based scRNA-Seq was carried out with Chromium Next GEN Single Cell 3’ Kit version 3 (10x Genomics), at the Eukaryotic Single Cell Genomics Facility at SciLifeLab, Sweden. The samples were processed with cellranger-4.0.0 pipeline (10x Genomics). The reads were mapped to a custom mouse (GRCm38) reference genome that contained GFP and Ai14 cassette (RFP and WPRE sequences) sequences. The reference genome was created with the 10x Genomics “cellranger mkref” and the mapping of the reads was done with the “cellranger count” function using default settings.

### scRNA-Seq analysis of adult lung cells

For the analysis of the droplet based scRNA-Seq dataset, we initially applied filtering criteria to filter out low quality cells and contaminants (Sftpc-GFP^pos^ library: GFP-UMIs>4, number of detected genes > 2500 and <5500 and percent of mitochondrial genes >0 and <7.5; Scgb1a1-CreER:Rosa-Ai14^pos^ Pdpn^neg^ library: RFP-UMIs>4, number of detected genes > 2500 and <5500 and percent of mitochondrial genes >0 and <7.5; Scgb1a1-CreER:Rosa-Ai14^pos^ Sftpc-GFP^pos^ library: RFP-UMIs>4 and GFP-UMIs>4, number of detected genes > 2500 and <5500 and percent of mitochondrial genes >0 and <7.5; Scgb1a1-CreER:Rosa-Ai14^pos^ Pdpn^pos^ library: RFP-UMIs>4, number of detected genes >3000 and <5500 and percent of mitochondrial genes >0 and <5). Genes with less than 50 counts in all cells were removed and the counts were transformed using the SCTransform^93^ function in Seurat^94^, with 4000 variable genes and regressing out the number of counts and detected genes and the percent of mitochondrial counts. The first 50 principal components were used for dimension reduction and clustering, setting the number of neighbours to 25 and the resolution to 0.2. MAST^95^ was used to identify DEGs after library normalization to 10.000 and log_2_-transformation.

We used an equal number of cells/cluster for the trajectory analysis and ran diffusion maps with Destiny^35^, implemented with scMEGA^96^. We used the 16 first principal components and k=25. We used the three first diffusion-map components for the visualization and down-stream analyses. We calculated the principal curves (“getCurves” function), the pseudotime estimates (“slingPseudotime” function) and the lineage assignment weights (“slingCurveWeights” function) with Slingshot^97^. We identified differentially expressed genes with the “fitGAM” function of tradeSeq^98^. For multiple trajectories, we used the “patternTest” and for one the “associationTest” functions. The genes were ordered based on the hierarchical clustering ward.D2 method, using “hclust” function in fastcluster package^99^ and plotted using a custom script. The “clusterboot” function of fpc package^100^ was used to calculate the stability values of gene-modules. GO-analyses were done at http://geneontology.org/ selecting as organism the Mus musculus and using default settings. The Fisher’s Exact test calculates the False Discovery Rate (FDR). Aggregated gene expression scores of genes in modules and biological processes were calculated with the “AddModuleScore” function in Seurat^94^. For Balloon-plots and heatmaps, we used the “DotPlot” and “DoHeatmap” functions in Seurat, in addition to the pheatmap-package^101^.

### Full length scRNA-Seq

Single-cell library preparation was done according to Smart-Seq2 protocol^65, 102^ with some modifications. Cells were sorted in 96-well plates (Piko PCR Plates 24-well, Thermo Scientific, SPL0240 and Plate Frame for 24-well PikoPCR Plates, Thermo Scientific, SFR0241). Each well contained Triton-X100 (0.2%) (Sigma-Aldrich, T9284-100ML), ERCC RNA Spike-In Mix (1:400.000) (Life-Technologies, 4456740), Oligo-dT30 VN (1.25µM) AAGCAGTGGTATCAACGCAGAGTAC(30 x T)VN, dNTPs (2.5mM/each) (Thermo Scientific-Fermentas, R0192) and Rnase Inhibitor (1U/µl) (Clontech, 2313A) in 4µl final volume. After sorting, strips were covered with Axygen PCR-tube caps (VWR, PCR-02-FCP-C), centrifuged and placed on dry ice until storage at −80°C for further use. For cell culture, cells were sorted into HBSS++ buffer and kept on ice until they were processed for culture. To optimize Smart-Seq2^65^ for mouse primary lung cells that are small and contain a small amount of RNA, we used 50% less Oligo-dT30 VN and the cDNA synthesis was divided into two steps, the first was without TSO LNA and the second contained 1µM TSO LNA and additional 40U of SuperScript II RT (Thermo-Fisher Scientific, 18064071). The reaction lasted 30 minutes at 42°C. Then, the enzyme was deactivated at 70°C for 15 minutes. For Pre-Amplification PCR, we used the KAPA HiFi Hotstart ReadyMix (2x) (KAPA Biosystems, KK2602) and the ISPCR-primer AAGCAGTGGTATCAACGCAGAGT. PCR included 21 cycles and the total volume increased to 50µl in order to reduce the concentration of the unused Oligo-dT30 VN and TSO-LNA primers.

Tagmentation and indexed library amplification were done with Nextera® XT DNA Library Preparation Kit (Illumina, FC-131-1096) and Nextera® XT Index Kit (96 indexes, 384 samples) (Illumina, FC-131-1002) according to the manufacturer protocol (with 2.5 x volume reduction in all reactions). For tagmentation, we used 50pg of the libraries, as it was indicated by the 500-9000bp fraction of the library (Bioanalyzer).

Sequencing was done with Illumina 2500 HiSeq Rapid mode using paired-end (2×125bp) and single-end (1×50bp) reading. For downstream analyses, we used one strand of paired-end libraries and trimmed the reads to 50bp.

### Single-cell RNA Sequencing bioinformatics analysis of Smart-seq2 dataset

We initially kept the libraries with >40% uniquely mapped reads to a reference genome that contained GFP and ERCC sequences and removed *Esr1*, as an artifact because of sequence similarities with Scgb1a1-CreER^T2^ transgene and *Xis*t. Individual sequencing datasets were filtered regarding the number of detected genes (lower threshold: 2000 genes and upper threshold 10000 (P2272, P2661) and 6000 (P3504, P7657)). Then, we filtered out the libraries with more than 200 counts of Pecam1 as not epithelial contaminants. Finally, we removed libraries with more than 7.5% of mitochondrial gene counts, resulting in 354 libraries for downstream analysis.

We used SCT-transformation in Seurat with 3000 variable genes and regressed out the number of counts and detected genes and the percent of mitochondrial counts. The 20 first principal components were used for dimension reduction, setting the number of neighbours to 12 and resolution to 1. Diffusion maps were produced as in the adult dataset using the first 12 principal components and k=12. For the identification of DEGs, we used the MAST analysis in Seurat. For the trajectory analyses, we used Slingshot, setting as root the cluster-0 (embryonic) and end-point clusters the −2 (S1), -1 (S2), -4 (ciliated) and -3 (DP). The diffusion-map 3D-plots were created with scatter3D function of scatterplot3d^103^. GO-analyses and aggregated scores were produced as in the adult dataset.

### scRNA-Seq of *Fgfr2*-inactivated airway epithelial cells

We followed the procedure for tissue dissociation and cell isolation as in the other FACs-sorting experiments. Single-cell suspensions from three P7 Scgb1a1-CreER^T2 neg/neg^; Rosa26-Ai14^pos/pos^; Fgfr2^fl/fl^ mice were pooled and used as negative control samples. Three P7 and Scgb1a1-CreER^T2 pos/neg^; Rosa26-Ai14^pos/pos^; Fgfr2^fl/fl^ mice were combined and used as experimental groups. The same approach was used for two P21 negative control lungs and three experimental. Cells were counted with a Biorad cell counter, blocked with TruStain FcX™ PLUS (Biolegend, 156604), stained with a PE/Cyanine7 anti-mouse CD326 antibody, and washed according to the manufacturer’s protocol (Biolegend, 118216). The negative control samples were sorted based on Epcam positivity and from the experimental groups, we isolated Epcam^pos^-Ai14^neg^ and Epcam^pos^-Ai14^pos^ cells. The isolated cells were processed with the Chromium Next GEM Single Cell 3ʹ Reagent Kits v3.1 (10xGenomics), following the manufacturer’s instructions and targeting 7000 cells/well. The produced libraries were sequenced with a NovaSeq 6000 in two runs, one for each time point.

### scRNA-Seq analysis of the *Fgfr2*-inactivated epithelial cells

We initially processed all cells and filtered out genes that were expressed in fewer than 5 cells and followed the same analysis approach as in the lineage-tracing dataset, using 4000 variable genes. We removed Krt13^high^ oesophageal/tracheal basal cells, Ptprc^pos^ immune and Col1a2^pos^ mesenchymal cells as contaminants. Filtered cells were re-clustered after filtering out genes that are expressed in less than 20 cells and selecting the 5000 most variable genes. We used the 20 first principal components, k=15 and resolution=0.2. Then, we selected the airway secretory clusters for downstream analyses as described for the other datasets, using 600 variable genes, 15 principal components, resolution = 0.99 and k=8.

### scRNA-Seq analysis of the GSE149563

For the analysis of the publicly available GSE149563 scRNA-Seq dataset, we analysed each timepoint individually with Seurat, using 4000 variable genes and 50 top principal components. We used DoubletFinder^104^ package in R to identify and remove multiplets. The postnatal datasets were integrated and processed for clustering and differential expression analysis as in the other datasets. The epithelial clusters were further filtered to remove possible endothelial (Pecam1^pos^) and mesenchymal (Col1a2^pos^) cells and re-clustered selecting the 4000 most variable genes. We used the 50 first principal components, k=25 and resolution=0.6.

### Organoid cultures

Lung digestion and cell sorting were performed as above, including the Dead cell stain (NucRed, Thermo) to sort out dead cells. Sorted epithelial cells (100-600 cells/well) were mixed with the Mlg2908 (ATCC, CCL-206) mouse lung fibroblasts (10^4^ cells/well), as described before^105^. Colonies were then fixed in 4% PFA O/N and placed in 30% sucrose solution for 24hours. Freeze-thawing and gentle pipetting were performed twice to remove Matrigel. Colonies were then incubated with 30% sucrose and 30% OCT overnight and embedded in OCT. Blocks were cut at 12-14µm for immunofluorescence.

### Immunofluorescence

The tissues were sectioned with a cryostat (Leica CM3050S). 10µm thick sections were placed on poly-lysine slides (Thermo Scientific, J2800AMNZ), kept at room temperature (RT) for 3hours with silica gel (Merck, 101969) to completely dry and then stored at −80°C until use. All antibodies are described in Supplementary Table 9. All secondary antibodies were used at a dilution of 1:300-1:400.

For antigen retrieval (when necessary, see Supplementary Table 9), slides were placed in plastic jars with the appropriate solution and warmed at 80°C for 30min in a water bath. Then, the jars were placed in ice for 30min to cool. Blocking was done with 5% donkey serum (Jackson Immuno-research, 017-000-121) for 1hour at RT and the primary antibodies were incubated at 4°C O/N. After washes, the secondary antibodies were applied on the sections at RT for 1hour in the dark. The nuclei were counterstained with a DAPI solution 0.5µg/µl (Biolegend, 422801) in PBS 1X Triton-X100 0.1% and for mounting we used the ProLong Gold Antifade Reagent (Thermo-Fischer Scientific, P36934).

In the staining for cells that escaped inactivation of Fgfr2, we used extended antigen retrieval incubation (90 minutes) and employed a Biotin-Streptavidin staining strategy to improve the FGFR2 signal. In short, we used the Avidin/Biotin Blocking Kit (SP-2001, Vector Laboratories) after blocking and before primary antibody incubation, according to the manufacturer’s suggestions (15min Avidin, Rinse, 15min Biotin, Rinse). After O/N incubation with primary antibodies and washes, we incubated the sections with a Biotin-SP-conjugated donkey anti-rabbit IgG (Secondary antibody) for 1hr at RT. After three washes, the sections were incubated with an Alexa Fluor® 647-conjugated Stretavidin for another 1hr at RT.

Images were acquired with Zeiss LSM780, LSM800 confocal microscopes (Carl Zeiss Microscopy GmbH, Jena, Germany) and Zeiss Axio Observer Z.2 fluorescent microscope with Colibri2 or Colibri7. Image analysis was done using Fiji^106^ and Zeiss Zen Blue 2.5.

### Quantification of S1 and S2 markers along the PD-axis

To quantify S1 or S2 marker co-expression, we acquired five confocal microscopy images from P, I1-3 and D domains, from one P60 mouse lung section. Cell counting was performed using a custom pipeline at Cell Profiler 3.1. “*Global*” threshold strategy, “*otsu*” threshold method and three classes of thresholding were used. False positive and negative cells were manually curated.

### Quantification of S1 and S2 markers in SPF and GF mice

For the quantification of mean fluorescence intensity of S1 or S2 marker in proximal and distal regions of the left lung lobe in germ-free (GF) (P62) and specific pathogen-free (SPF) mice (P60), z-stacks of the whole lobe were captured with a Zeiss Axio Observer Z.2 fluorescent microscope with Colibri2 or Colibri7, equipped with a Zeiss AxioCam 506 Mono digital camera and an automated stage. Z-Stacks were projected using the “Orthogonal Projection” using the “Maximum” method and stitched using the “Stitching function” (Zen Blue). ROIs were drawn manually in proximal and TB regions using E-Cadherin and DAPI channels as a reference, and mean fluorescence intensity was measured using the Zen Blue software. Mean Fluorescence intensity (MFI) for the individual markers was normalized against the MFI of E-Cadherin for each ROI and the results from 3 animals per condition (SPF or GF) were combined into one dataset. Statistical analysis for differences between proximal and distal was done using a two-way unpaired T-test in GraphPad Prism.

### SCRINSHOT spatial analyses

For spatial analysis of the identified cell types, we applied SCRINSHOT^57^. The utilized padlock and detection probes are summarized in Supplementary Table 17. Images were captured with a Zeiss Axio Observer Z.2 fluorescent microscope with Colibri2 or Colibri7, equipped with a Zeiss AxioCam 506 Mono digital camera and an automated stage.

We used DAPI to align the images of the same areas between the hybridizations. We created multi-channel *.czi files with the signal of each detected gene as a unique channel and exported them as images (16-bit *.tiff format) using Zen Blue 2.5 (Carl Zeiss Microscopy, GmbH). Images were tiled with Matlab with Image Analysis toolbox (The MathWorks, Inc.). Manual nuclear segmentation was done with Fiji ROI Manager^106^ and signal-dot counting was performed with Cell-Profiler 3.15^107^. Annotation of signal dots to the cells (2µm expanded nuclei) was done with Fiji.

### S1 and S2 cell spatial analyses

For the spatial analysis of S1- and S2-cells, we targeted the module-5 secreted proteins *Scgb3a1, Reg3g*, *Bpifb1*, *Tff2* and *Muc5b*, the receptor *Lgr6* that has previously been reported to be expressed by distinct epithelial and mesenchymal lung cell-types^108^ and the goblet-cell transcription factor *Pax9*, which are enriched in S1-cells. For the distal lung compartment (DP, S2 and alveolar cells), we used the module-2 markers *Hp* and *Scgb1a1* in addition to the module-3 surfactant proteins *Sftpc* and *Sftpb*, the enzyme *Atp8a1*, the IGF-signaling regulator *Igbp6* and the advanced glycosylation end products receptor *Ager*. We also targeted the ciliated-cell markers *Foxj1* and *Tuba1a* to recognize ciliated cells.

P, I1, I2, I3 and D domains from sections of three P60 mouse lungs were analysed for the selected panel of markers. Nuclei were manually segmented in the acquired images based on DAPI and manually curated based on E-cadherin antibody staining, resulting to 6915 nuclei. Nuclear ROIs for each animal were expanded and filtered, keeping those with size between Mean cell-ROI size ± 2 standard deviations. Cell-ROIs with dots for only 1 analysed marker were removed. We further filtered the Cell-ROIs, keeping those with a total number of dots between the Mean number of dots ± 2 standard deviations. Cell-ROIs from all images were merged and log_2_-transformed [log_2_(dots + 1)]. After principal component analysis (PCA), the top up- or down-regulated genes of the first two principal components were used to cluster the Cell-ROIs with clusterboot, using the ward.D2 method. The heatmap of the analysed cells was done with pheatmap-package in R. The balloon-plots of the expression levels (color intensity) and the percent of positive cells (size) were produced with ggpubr-package in R.

### DP cell spatial analysis

For the selection of the gene-panel for the spatial analysis of DP-cells, we used differential expression analyses to compare them with S2-cells and AEC2a. The panel included *Scgb1a1* and *Sftpc* (the positivity of both defines the DP-cells/BASCs^22, 24, 25^) and the cytochrome genes *Cyp2f2* and *Cyp4b1* that showed high expression in S1- and S2-cells, moderate in DP and low in AEC2 cells. Based on the average fold change and the percentage of positive cells, we additionally selected the secreted proteins *Lyz2* and *Lgi3*, the extracellular matrix proteins *Egfl6*, *Npnt* and *Col4a2*, the enzyme *Napsa*, the surface molecules *Cd74* and *Cldn18*, the transcription factors *Etv5* and *Rbpjl* and the negative regulator of Wnt-signalling *Nkd1*. We also included *Ager* as a distal epithelial marker and the NE-cell markers, *Ascl1* and *Calca* (Cgrp). In two independent experiments, we analysed several lung areas from three adult (P60) mice and manually segmented 58072 nuclei. Cell-ROIs with a size outside 2 standard deviations from the average size of all Cell-ROIs from each lung were excluded from the analysis. The selection of ROIs extracted DP-cells with more than 24 dots of *Scgb1a1* and *Sftpc* that have also *Scgb1a1*-dots ≥ 10 and *Sftpc*-dots ≥ 10. The balloon-plots of the expression levels (colour intensity) and the percent of positive Cell-ROIs (size) were produced with ggpubr-package^109^ in R.

### Statistical analyses

Statistical analysis of the results was done with a two-way multiple comparisons test in GraphPad Prism (GraphPad Software, Inc.) or by MAST in Seurat. In GraphPad Prism, multiple comparisons were performed using Tukey statistical hypothesis testing. Adjusted p-values in MAST were calculated based on Bonferroni correction.

## Data Availability

scRNA-Seq data are available in GEO (lineage-tracing dataset of Scgb1a1-CreER^T2pos/neg^;Rosa26-fGFP^pos/neg^ cells: GSE215957, adult dataset of Scgb1a1-CreER^T2 pos/neg^;Rosa26-Ai14^pos/neg^ cells: GSE216210 and Fgfr2-inactivation dataset of Scgb1a1-CreER^T2 pos/neg^;Rosa26-Ai14^pos/pos^; Fgfr2^fl/fl^ cells: GSE216451). Scripts and RAW-image data can be found in Zenodo (doi: 10.5281/zenodo.10418253).

## Acknowledgements

We thank the SciLifeLab NGI and WABI for long-term bioinformatics support. We acknowledge resources provided by the Swedish National Infrastructure for Computing (SNIC) at UPPMAX, partially funded by the Swedish Research Council through grant agreement no. 2018-05973 (projects b2015134 and SNIC 2021/22-431). The work was supported by grants from VR 2019-04893, CF 211794 Pjo1H and the Erling Persson Foundation 2023-0035 to CS. We acknowledge the European Respiratory Society-EMBO for the European Respiratory Society Long-Term Research Fellowship to AS (Reference Number: LTRF 2014 – 3565).

---

**Extended Data Figure 1.**
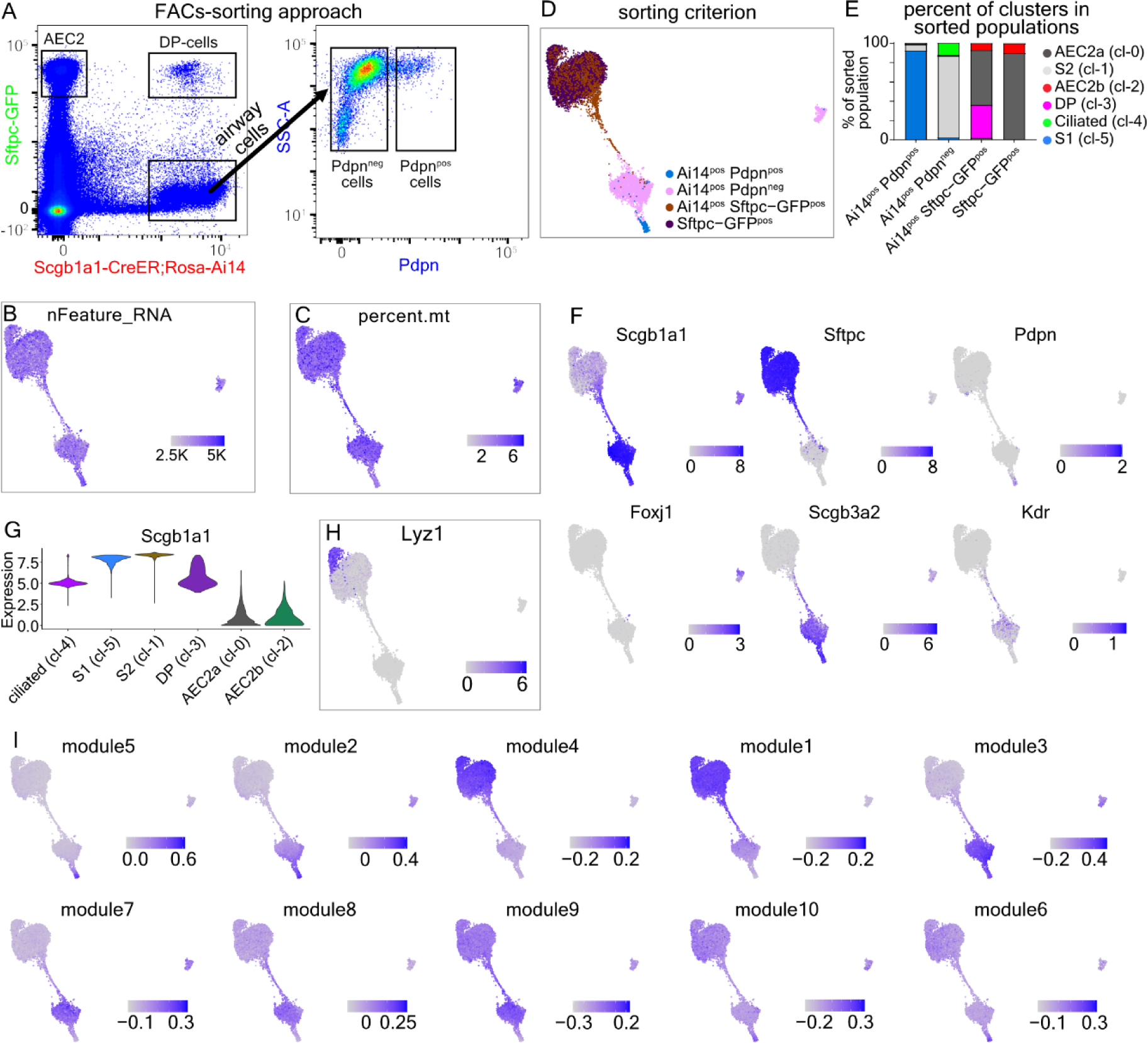
Characterization of lung secretory cell heterogeneity. **(A)** FACS-sorting approach for isolation of AEC2, DP-cells and S1- with S2-cells (left) and S2 (Pdpn^neg^) and S1 (Pdpn^pos^) cells (right). **(B-C)** UMAP-plots of showing numbers of detected genes and the percentage of mitochondrial genes, respectively. **(D)** UMAP-plot showing the corresponsdance of sorting criteria for cell isolation for each cluster. **(E)** Bar-plot showing the percentage of cells in the clusters, according to the sorting criteria. **(G)** Violin plot of *Scgb1a1* expression in the clusters. **(H)** UMAP-plot of Lyz1 expression. **(F)** UMAP-plots of the known cell-type markers *Scgb1a1, Sftpc, Pdpn, Foxj1, Scgb3a2* and *Kdr*. In “G-F”, expression levels as log_2_(normalized UMI-counts+1) (library size was normalized to 10.000). **(I)** UMAP-plots showing the aggregated expression scores of the genes in the 10 suggested modules (Fig. 2B). Blue: high, Gray: low.

**Extended Data Figure 2.**
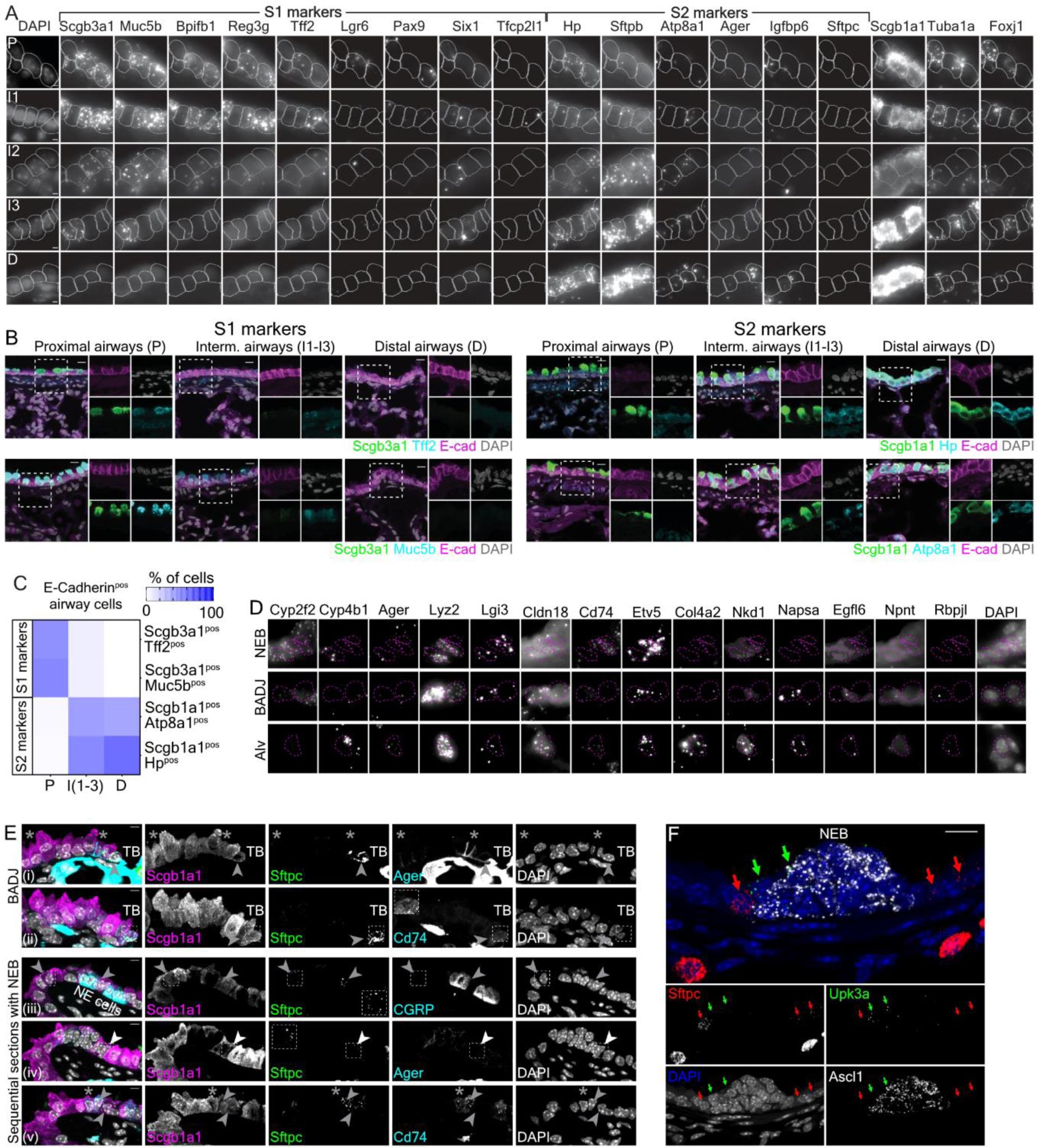
Spatial validation of gene expression markers. **(A)** Characteristic examples of detected mRNA molecules with SCRINSHOT for the analysed genes, in the indicated domains. Projected cell-ROIs: manually segmented nuclei, expanded for 2 µm. **(B)** Confocal microscopy images of immunofluorescence for: Scgb3a1, E-cadherin (Ecad) and Tff2 or Scgb3a1, Ecad and Muc5b in proximal (left), intermediate (middle) and distal (right) airway regions. Scgb3a1: green, Muc5b or Tff2: cyan, Ecad: magenta, Nuclei (DAPI): grey. (Bottom-S2 markers) Scgb1a1, Ecad and Hp or Scgb1a1, Ecad and Atp8a1. Scgb1a1: green, Hp or Atp8a1: cyan, Ecad: magenta, Nuclei (DAPI): grey. Scale bar:10 µm. **(C)** Heatmap showing the percent of double positive cells of the indicated markers, in the proximal, intermediate and distal airways. Five images/domain were quantified. Scgb3a1-Tff2 staining: P: 141 cells, I (1-3): 157 cells, D: 209 cells, Scgb3a1-Muc5b staining: P: 149 cells, I (1-3): 194 cells, D: 181 cells, Scgb1a1-Hp staining: P: 123 cells, I (1-3): 167 cells, D: 170 cells, Scgb1a1-Atp8a1 staining: P: 120 cells, I (1-3): 186 cells, D: 172 cells. **(D)** Single-channel images of the analysed ROIs in Fig. 3F, showing SCRINSHOT signal. **(E)** Representative confocal images of Scgb1a1 and Sftpc immunofluorescence in combination with Ager (i) and Cd74 (ii) in TBs of an adult lung section. Arrows: Scgb1a1^pos^ Sftpc^pos^ cells and asterisks: Scgb1a1^pos^Ager^pos^Sftpc^pos^ cells, showing that airway epithelial cells that expressed Sftpc and/or Cd74 and vice versa, suggesting for additional heterogeneity within airway epithelium (iii-v) Sequential section images of the same airway junction showing Scgb1a1^pos^Sftpc^pos^ cells in relation to CGRP (iii), Ager (iv) and Cd74 (v). (iii-iv) Arrows: Scgb1a1^pos^Sftpc^pos^ cells. (v) Arrow: a Scgb1a1^pos^SftpcposCd74^pos^ cell, asterisk: a Scgb1a1^pos^SftpcnegCd74^pos^ and arrowhead: a Scgb1a1^pos^SftpcnegCd74^neg^. Scgb1a1^pos^ Sftpc^pos^ Ager^pos^ cells are only found in TBs suggesting that DP-cells are only a subset of the very distal airway Scgb1a1^pos^ Ager^pos^ cells. Images are maximal-intensity projections of 9 z-stacks. Inserts: magnifications of the indicated areas. Scale-bar 5 µm. “DP”: Scgb1a1^pos^ Sftpc^pos^ cells. **(F)** Confocal image of a representative neuroepithelial body (NEB), showing that Upk3a^pos^ cells are mainly found over the neuroendocrine Ascl1^pos^ cells, whereas Sftpc^pos^ airway cells are localized laterally of them. Sftpc: red, Upk3a: green, Ascl1: grey and DAPI: blue. Scale bars 10 µm. “PD”: proximal-distal, “DP”: Scgb1a1^pos^ Sftpc^pos^ double-positive. “Alv”: alveolar, “Arw”: airway, “TB”: terminal-bronchiole, “BADJ”: bronchiole-alveolar duct junction. At least three lungs were analysed for each experiment with similar results.

**Extended Data Figure 3.**
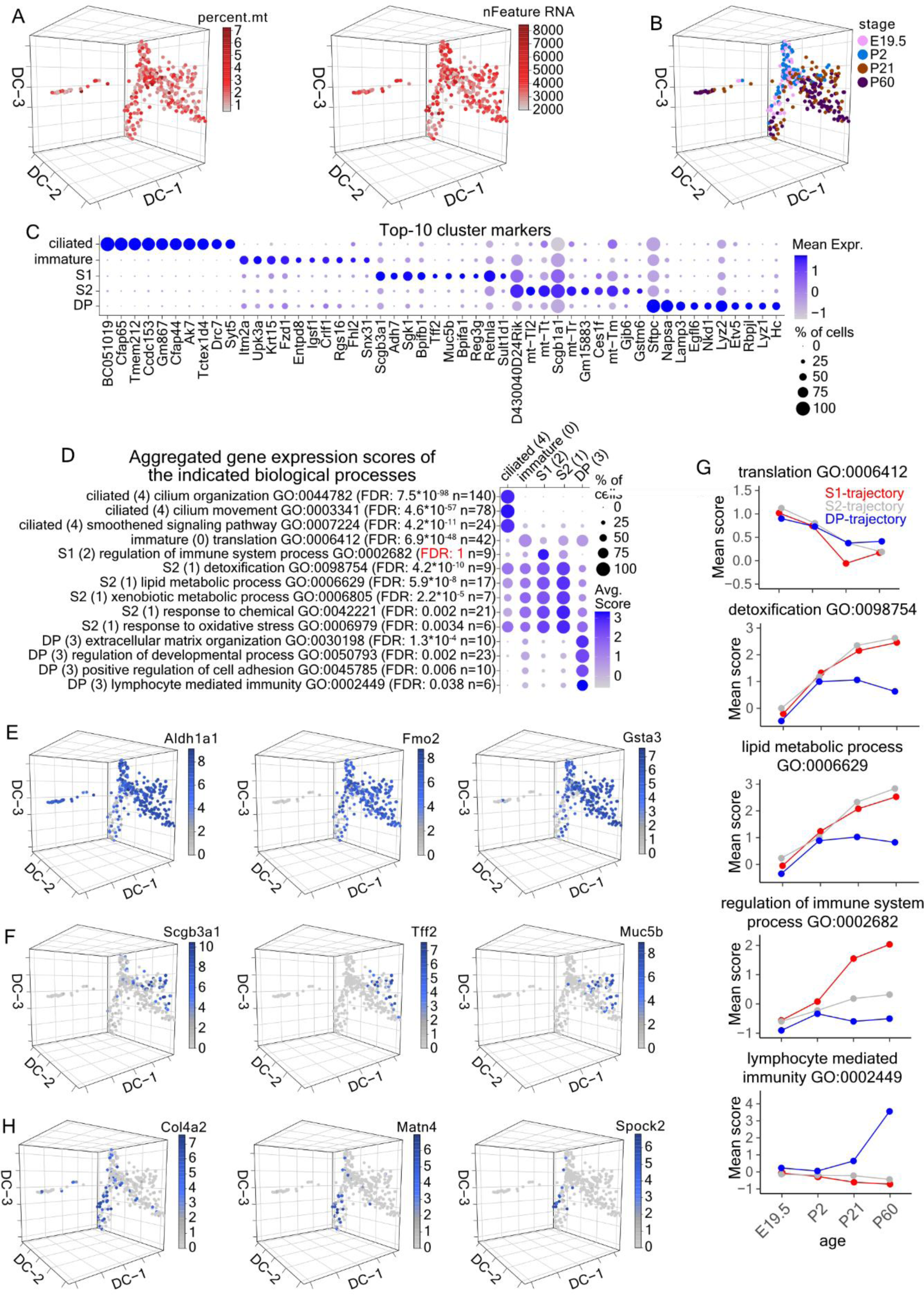
Lineage-tracing of airway secretory cell heterogeneity. **(A)** 3D diffusion-map plots of 354 full-length, single-cell cDNA libraries, showing the percent of mitochondrial counts (left) and number of detected genes (right). Red: high, Gray: low. **(B)** Diffusion-map plot showing the stage of the analysed cells. **(C)** Balloon-plot of the top-10 cluster markers, according to MAST differential expression analysis. The genes were filtered according to average log2 Fold-change (>0.5), adjusted p-value (<0.05) and percent of positive cells (>0.25) and the top-10 markers according to average log2 Fold-change were plotted. Gene order follows the cluster order. Balloon size: percent of positive cells. Colour intensity: scaled expression. Blue: high, Gray: low. **(D)** Balloon-plot of the average, aggregated gene expression scores of selected biological processes. The analysis is based on the statistically significant cluster markers (Suppl. Table 6). Balloon size: percentage of positive cells. Colour intensity: aggregated expression. Blue: high, Gray: low. “FDR”: false discovery rate, “n”: number of genes. **(E-F)** Diffusion-map plots showing the expression of the metabolic enzymes *Aldh1a1*, *Fmo2* and *Gsta3* (E) and of the innate immunity genes *Scgb3a1*, *Tff2* and *Muc5b* (F). Expression levels: log_2_(normalized counts+1) (library size was normalized to 10^6^). Blue: high, Gray: zero. **(G)** Line-plots of the Mean aggregated gene expression scores of the indicated biological processes in the S1 (left) and S2 (right) cell clusters, according to their age. **(H)** As in “D” for the extracellular matrix proteins *Col4a2*, *Matn4* and *Spock2* that are expressed in the middle part of the DP-trajectory.

**Extended Data Figure 4.**
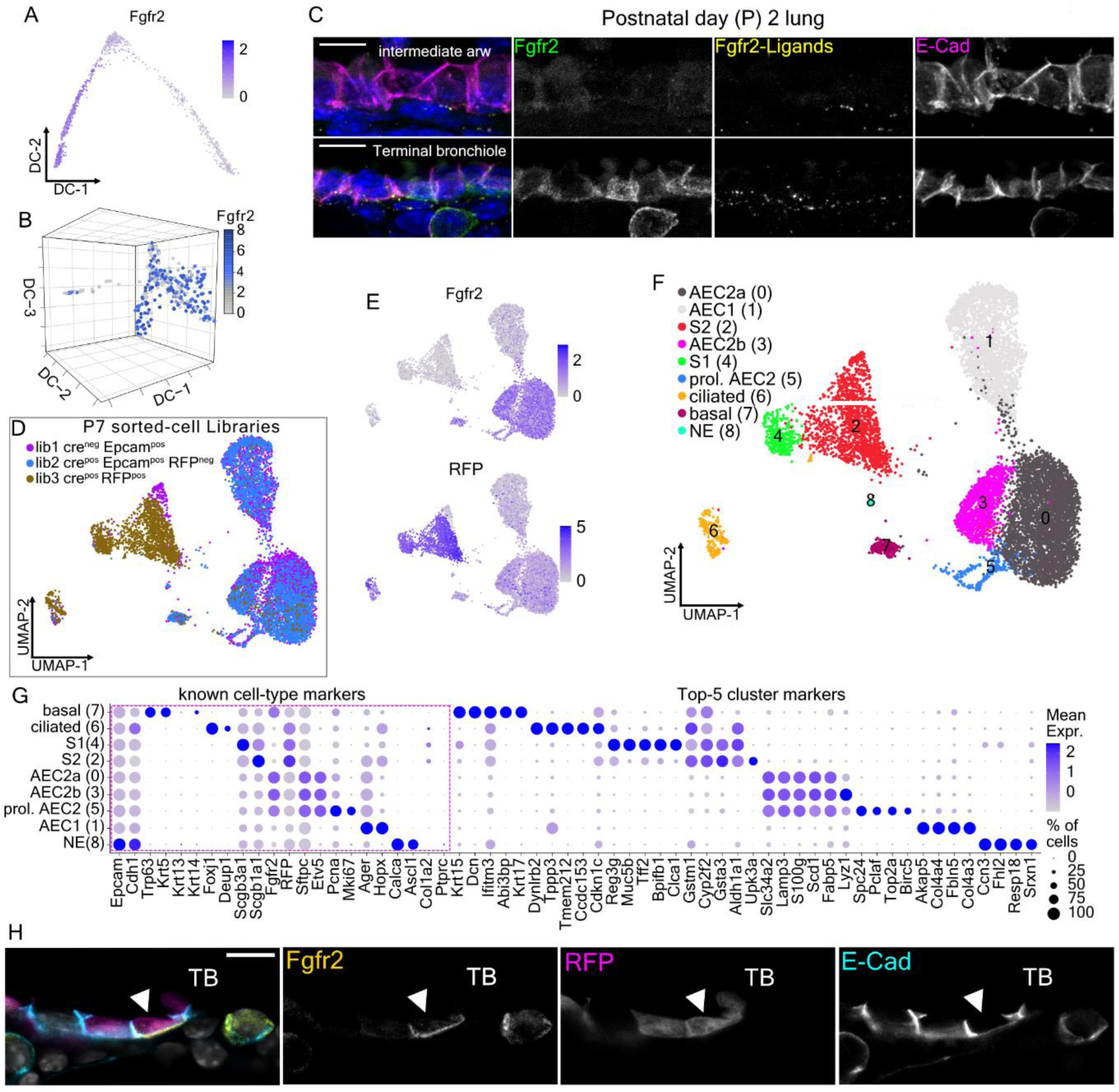
*Fgfr2* expression and its inactivation in airway epithelium. **(A-B)** Diffusion-map plots showing *Fgfr2* expression in the adult (top) and lineage tracing (bottom) scRNA-Seq datasets. Expression levels: (A) log_2_(normalized UMI-counts+1) (library size was normalized to 10.000), (B) log_2_(counts+1) (library size was normalized to 10^6^). **(C)** Confocal microscopy z-stack projection images of representative intermediate (top) and distal (bottom) airways for E-cadherin (red), Fgfr2 (green), Fgfr2 ligands (Fgfr2β (IIIb) Fc chimeric protein, yellow) and nuclei (DAPI-blue). Images are maximal-intensity projections of 8 z-stacks. Scale bar: 10µm. **(D)** UMAP-plot showing the sequenced library-information of the analysed cells. **(E)** UMAP-plots showing the Fgfr2 (top) and RFP (bottom) expression in the analysed dataset. Expression levels as log_2_(normalized UMI-counts+1) (library size was normalized to 10.000). **(F)** UMAP-plot showing the cell clusters and their annotations. **(G)** Balloon-plot of known cell-type markers (*Epcam-Ptprc*) and of the top-5 cluster markers. Genes were filtered according to average adjusted p-value (<0.05), percent of positive cells in the corresponding cluster (>0.25) and difference in positive cells (>0.5) and the top-5 markers according to average log2 Fold-change were plotted. Gene order follows the cluster order. Balloon size: percent of positive cells. Colour intensity: scaled expression. In all plots, blue: high, grey: zero/low. **(H)** Single-plane confocal microscopy images of a terminal bronchiole from a postnatal day-7 (P7) Fgfr2-mutant lung. Immunofluorescence for Fgfr2 (yellow), Rosa26-Ai14 (red), E-cadherin (cyan) and nuclei (DAPI-grey). Arrow-head: airway epithelial that recombined the Rosa26-RFP locus but remains Fgfr2^pos^, indicating inefficient Fgfr2-inactivation. Scale-bar 10µm.

**Extended Data Figure 5.**
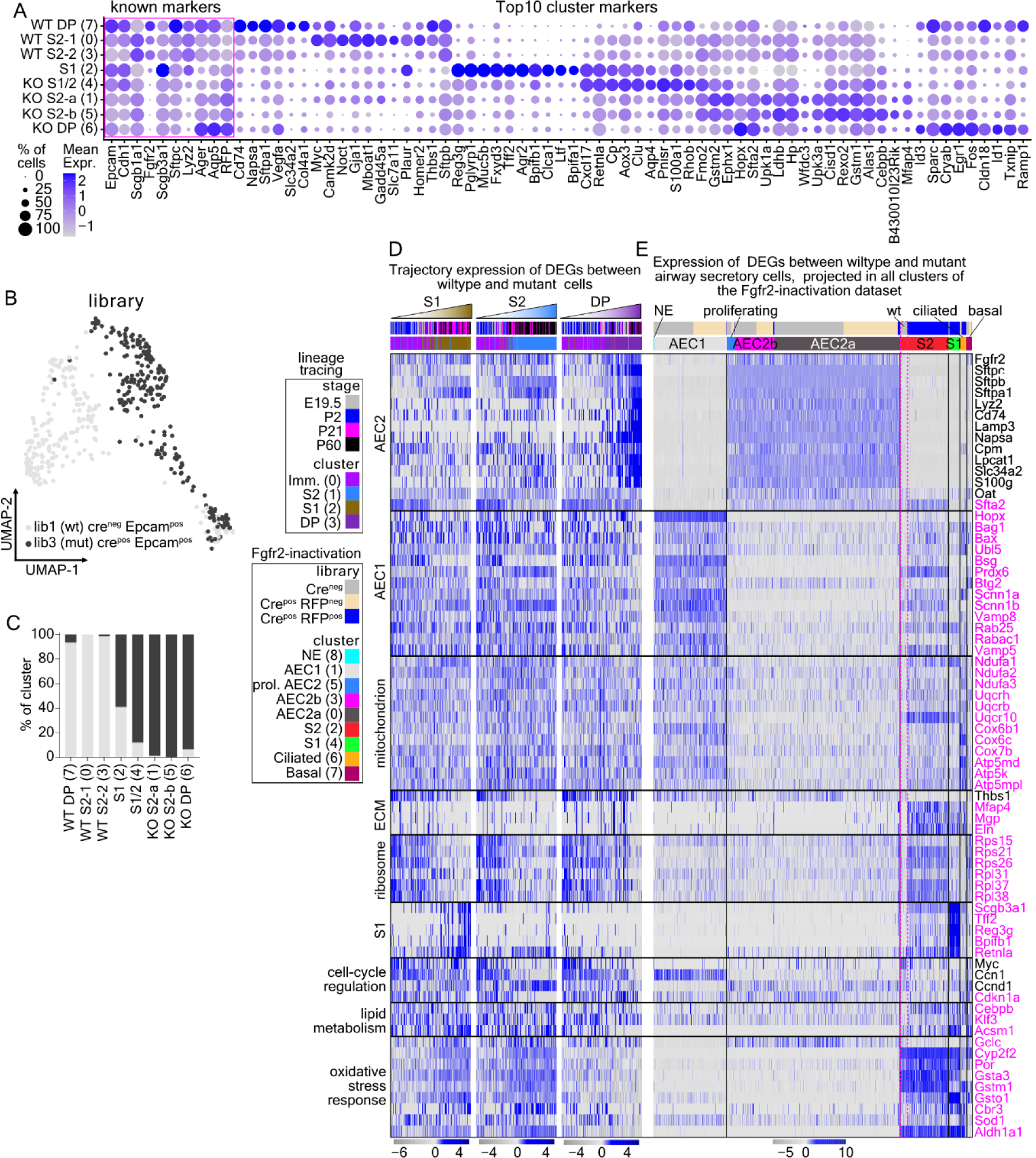
*Fgfr2* inactivation in airway epithelium. **(A)** Balloon-plot of known cell-type markers (*Epcam-RFP*) and of the top-10 cluster markers. Genes were filtered according to adjusted p-value (<0.05), percent of positive cells in the corresponding cluster (>0.5) and difference in positive cells (>0.25). The top-20 markers were selected according to average log2 Fold-change and the top-10 were plotted according to the difference in positive cells. Gene order follows the cluster order. Balloon size: percent of positive cells. Colour intensity: scaled expression. Blue: high, Gray: low. **(B)** UMAP-plot of the analysed airway secretory cells, showing the library-information. Light-grey: wildtype library-1. Dark-grey: mutant library-3. **(C)** Bar-plot of the percent of the two libraries in each cluster. Colours as in “B”. **(D)** Heatmap of the lineage-tracing dataset (see Fig. 5B), showing the expression of selected, differentially expressed genes between the wildtype (library-1) and the mutant (library-3) airway secretory cells. **(E)** As in “D” for the and whole Fgfr2-inactivation dataset (see Extended Data Fig. 4F). Gene-order as in Fig 6C. Colour: scaled expression. blue: high, grey: low.

**Extended Data Figure 6.**
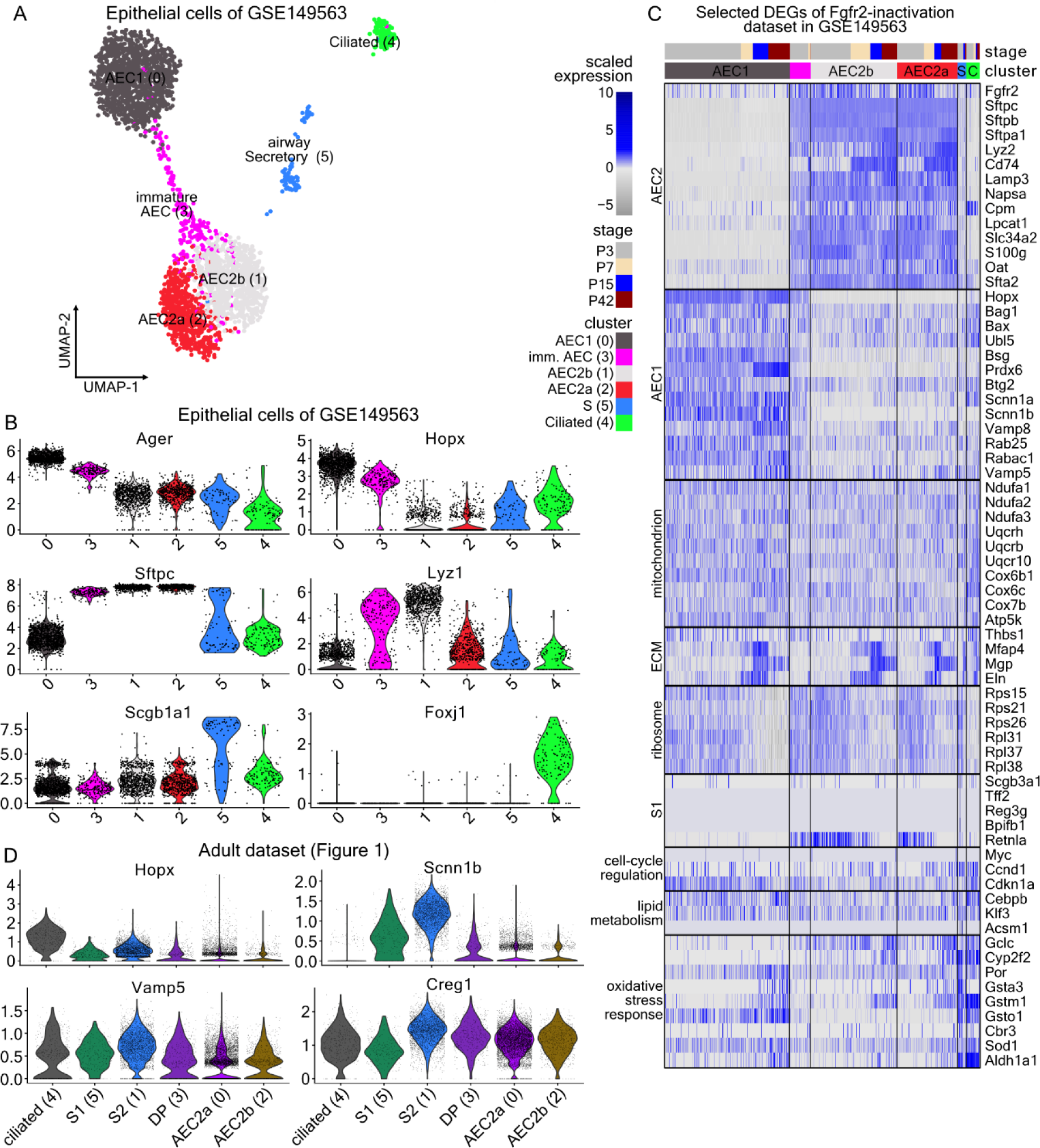
Epithelial cell analysis of the GSE149563. **(A)** UMAP-plot of 3053 Pecam1^neg^ Col1a2^neg^ epithelial cells of the publicly available GSE149563 single-cell RNA Sequencing dataset. Colours: suggested clusters. **(B)** Violin-plots of known cell-type markers that were used for cluster-annotations. *Ager* and *Hopx* (AEC1), *Sftpc* and *Lyz1* (AEC2), *Scgb1a1* (airway secretory) and *Foxj1* (ciliated). **(C)** Heatmap of the epithelial cells of GSE149563, ordered by cluster, showing the expression of selected, differentially expressed genes between the wildtype (library-1) and the mutant (library-3) airway secretory cells. Gene-order as in Fig. 6C. Colour: scaled expression. blue: high, grey: low. **(D)** Violin-plots of *Hopx*, *Scnn1b*, *Vamp5* and *Creg1* in the adult cell dataset of the present study (colours as in Fig. 1C). In all violin-plots expression levels as log_2_(normalized UMI-counts+1) (library size was normalized to 10.000).

**Supplementary Figure 1.**
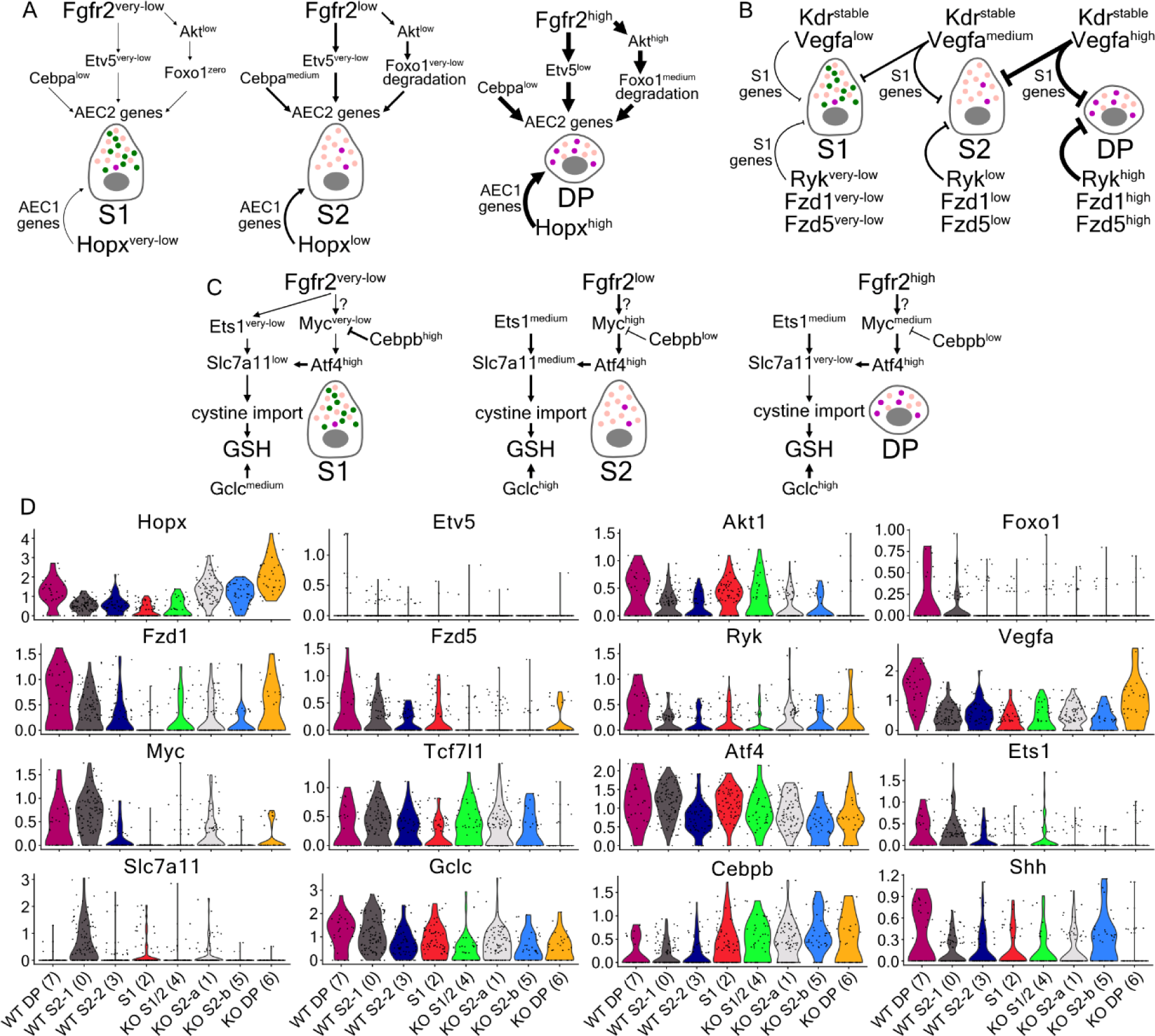
Supporting information to Figure 7 proposed models. **(A-C)** Proposed models of signaling pathways in the normal epithelium, as in Figure 7. **(D)** Violin plots showing the expression levels of the included genes in the models that change upon Fgfr2-inactivation. Expression levels: log_2_(normalized UMI-counts+1) (library size was normalized to 10.000).

**Supplementary Table 1.** Results of the differential expression analyses of the adult single-cell RNA Sequencing dataset with MAST. **(A)** Comparison of each cluster against all others. **(B)** Filtered genes of “A”, adjusted p-value <0.05, pct.1 (percent of positive cells in the corresponding cluster) >0.25, log2 Fold-change >0.5. **(C)** Comparison of the two alveolar epithelial cell type 2 (AEC2) clusters −0 and −2. **(D)** Comparison of the airway secretory clusters −1 (S2) and −5 (S1). **(E)** Comparison of the AEC2a cluster-0 with the double positive (DP) cluster-3. **(F)** Comparison of the airway S2 cluster-1 with the double positive (DP) cluster-3.

**Supplementary Table 2.** Results of the gene ontology analyses. The tables include the enriched biological processes in each of the adult cell clusters, based on the genes in Suppl. Table 1B. The results were filtered for false discovery rate (FDR) <0.05 and fold enrichment >2.

**Supplementary Table 3**. Differentially expressed genes and their related biological processes along pseudotime trajectory. **(A)** The 10 modules of differentially expressed genes along the adult secretory cell pseudotime trajectory. **(B-G)** Results of the gene ontology analyses, showing the enriched biological processes of each gene-module. The modules 8-10 did not produce enriched biological processes with false discovery rate (FDR) <0.05 and fold enrichment >2. **(H-I)** Detailed information of the biological processes and genes in Fig. 2C-D.

**Supplementary Table 4.** Positional statistics of double positive (DP) Scgb1a1^pos^ Sftpc^pos^ cells using one-way ANOVA with Kruskal-Wallis multiple comparisons test.

**Supplementary Table 5.** Results of the differential expression analysis of all clusters in the lineage-tracing dataset, using MAST. **(A)** Comparison of each cluster against all others. **(B)** Filtered genes of “A” with adjusted p-value <0.05, **(C)** Filtered genes of “A” with adjusted p-value <0.05, pct.1 (percent of positive cells in the corresponding cluster) >0.25, log2 Fold-change >0.5.

**Supplementary Table 6.** Results of the gene ontology analyses. The tables include the enriched biological processes in each of the lineage-tracing dataset cell clusters, based on the genes in Suppl. Table 5B. The results were filtered for false discovery rate (FDR) <0.05 and fold enrichment >2.

**Supplementary Table 7.** Results of the differential expression analyses from the Fgfr2-inactivation experiment with MAST. **(A)** Comparison of each cluster against all others from the whole Fgfr2-inactivation dataset (see Extended Data Fig. 4F). **(B)** Comparison of each cluster against all others from the airway secretory cell dataset (see Fig. 6B). **(C)** Filtered genes of “B”, using adjusted p-value <0.05, percent of positive cells in the corresponding cluster (>0.5) and log2 Fold-change >0.25. **(D)** Comparison of the wildtype (library-1) with the mutant (library-3) cells of the airway secretory cell dataset (see Extended Data Fig. 5B). **(E)** Filtered genes of “D”, using adjusted p-value <0.05, log2 Fold-change >0.25. **(G)** Genes of “E” including the gene description, gene type, how it changes compared to wildtype cells and the most relevant biologicals process, molecular functions and cellular components retrieved from Biomart database of ensembl.org. Additional information about the gene function was also included if available. Red-font: selected genes for heatmap plots in Fig. 6C and Extended Data Fig. 5D-E and 6C. **(G)** Transcription Factors that are found in “C”.

**Supplementary Table 8.** Results of the gene ontology analyses. The tables include the enriched cellular components according to the differentially expressed genes (see Suppl. Table 7E) of airway secretory **(A)** wildtype (library-1) and **(B)** mutant cells (library-3). The results were filtered for false discovery rate (FDR) <0.05 and fold enrichment >2. **(C)** Information about the terms in Fig. 6D.

**Supplementary Table 9.** List of the used antibodies in the study.

**Supplementary Table 10.** List of used SCRINSHOT probes in the study.

